# ERAP1 activity modulates the immunopeptidome but also affects the proteome, metabolism and stress responses in cancer cells

**DOI:** 10.1101/2024.09.20.613815

**Authors:** Martha Nikopaschou, Martina Samiotaki, Ellie-Anna Stylianaki, Kamila Król, Paula Gragera, Aroosha Raja, Vassilis Aidinis, Angeliki Chroni, Doriana Fruci, George Panayotou, Efstratios Stratikos

**Affiliations:** National Centre for Scientific Research Demokritos, Agia Paraskevi, Greece; Department of Chemistry, National and Kapodistrian University of Athens, 15784 Zografou, Greece; Biomedical Sciences Research Center “Alexander Fleming”, Institute for Bioinnovation, 16672 Vari, Greece; Biomedical Sciences Research Center “Alexander Fleming”, Institute for Fundamental Biomedical Research, Vari, Greece; Division of Pediatric Hematology and Oncology, Bambino Gesù Children’s Hospital, IRCCS, Rome, Italy; Center for Translational Immunology, University Medical Center Utrecht, Utrecht University, Utrecht, The Netherlands

## Abstract

Endoplasmic reticulum (ER) aminopeptidase 1 (ERAP1) metabolizes peptides inside the ER and shapes the peptide repertoire available for binding to Major Histocompatibility Complex Class I molecules (MHC-I). Moreover, it may have additional effects on cellular homeostasis, which have not been explored. To address these questions, we used both genetic silencing of ERAP1 expression as well as treatment with a selective allosteric ERAP1 inhibitor to probe changes in the immunopeptidome and proteome of the A375 melanoma cancer cell line. We observed significant immunopeptidome shifts with both methods of functional ERAP1 disruption, which were distinct for each method. Both methods of inhibition led to an enhancement, albeit slight, in tumor cell killing by stimulated human PBMCs and in significant proteomic alterations in pathways related to metabolism and cellular stress. Similar proteomic changes were also observed in the leukemia cell line THP-1. Biochemical analyses suggested that ERAP1 inhibition affected sensitivity to ER stress, reactive oxygen species production and mitochondrial metabolism. Although the proteomics shifts were significant, their potential in shaping immunopeptidome shifts was limited since only 15.8% of differentially presented peptides belonged to proteins with altered expression and only 5.0% of proteins with altered expression were represented in the immunopeptidome shifts. Taken together, our findings suggest that modulation of ERAP1 activity can generate unique immunopeptidomes, mainly due to altered peptide processing in the ER, but also induce changes in the cellular proteome and metabolic state which may have further effects on tumor cells.

## INTRODUCTION

The ability of viruses and tumors to hide within autologous cells poses a major challenge for the immune system. To overcome this challenge, the immune system has developed the major histocompatibility complex class I (MHC-I) antigen presentation pathway, which is active in almost all cell types ^1,2^. In this pathway, MHC-I molecules bind fragments of intracellular proteins and transport them to the cell surface ^2,3^. There, the peptide/MHC-I complexes are presented to cytotoxic T lymphocytes, which can distinguish between native and foreign proteins and kill cells expressing viral proteins or tumor antigens^1–5^. Natural killer cell responses can also be regulated by MHC-I cargo and complement these effects^6^. For effective immune surveillance, a broad peptide repertoire is typically presented^5^. This repertoire is collectively known as the immunopeptidome^7^.

The generation of the immunopeptidome usually begins with the degradation of intracellular, often defective^8^, proteins by the proteasome. The peptide products of this degradation are delivered to the endoplasmic reticulum (ER) via the Transporter Associated with Antigen Processing (TAP)^9^. Peptides generated by the proteasome are between 3 and 22 amino acids long, while only about 15% correspond to the optimal length for MHC class I binding (8-9 amino acids). Another 15% are longer than 10 amino acids and could serve as peptide precursors for loading on MHC-I^10^. These precursors can further be trimmed to the optimal length by the ER-resident aminopeptidases ERAP1 and ERAP2^11–14^. Trimmed peptides can be loaded onto MHC-I molecules with the help of the Peptide Loading Complex (PLC), a multi-protein machinery consisting of the MHC-I together with TAP, tapasin, ERp57 and calreticulin^15^. The peptide-loaded MHC-I are finally transported to the cell surface via the Golgi complex^4,9^.

Given the importance of the immunopeptidome in adaptive immunity, several efforts are underway to better understand how it is generated and how it can be modulated for therapeutic applications in the context of cancer, autoimmunity, and infections^7^. Amongst others, modulation of the immunopeptidome can be achieved by perturbations in the activity or expression of proteins in the antigen processing and presentation pathway^16^, including ERAP1 and its murine homologue ERAAP^17–20^.

Modulation of ERAP1 activity can be induced *in vitro* in two different ways: genetic knock-out of the enzyme or pharmacological modulation. The development of ERAP1 inhibitors has been ongoing for several years, and many of the efforts have focused on the active site of the enzyme which contains a zinc atom^21,22^. However, as the active site is largely conserved amongst other family members, active-site inhibitors often suffer from limited selectivity^23^. An alternative approach that has been gaining traction recently is the targeting of ERAP1 allosteric sites with compounds that display enhanced selectivity ^23,24^.

In addition to MHC-I antigen processing and presentation, ERAP1 has been suggested to participate in other biological functions, including blood pressure regulation^25,26^, angiogenesis^27^, ectodomain shedding of cytokine receptors^28,29^, Hedgehog-dependent tumorigenesis^30^, innate immunity^31,32^ and ER stress^33^. Since indirect proteomic alterations can contribute to changes in the immunopeptidome^34^, ERAP1 could also affect the immunopeptidome indirectly, through its effect in some of the aforementioned functions. At the same time, potential effects of ERAP1 in pathways other than antigen processing and presentation may also have therapeutic potential. Consequently, it is essential to simultaneously evaluate the effects of ERAP1 inhibition on the immunopeptidome and overall cellular functions. This can be performed in an unbiased way by evaluating the changes in the cellular proteome after ERAP1 modulation, as proteins are effectors of biological function^35^.

To better understand the potential interplay between direct and indirect regulation of the immunopeptidome by ERAP1, as well as explore its role in cellular homeostasis we combined immunopeptidomic with proteomic and biochemical analyses in the melanoma cell line A375, a well-established cellular system for exploring immunopeptidome modulation^18^. We used a data-independent acquisition (DIA) strategy to analyze how the immunopeptidome and proteome of A375 cells are altered after genetic or pharmacological ERAP1 inhibition, utilizing a recently developed allosteric inhibitor^24^. We demonstrate that although ERAP1 activity is critical for immunopeptidome regulation, its function also impacts cellular homeostasis, and its activity can induce changes related to mitochondrial metabolism and ER stress. Our data suggest that ERAP1 activity, while highly specialized for antigen presentation, is also important for ER peptide homeostasis and may therefore play a dual role in metabolic adaptations of tumor cells attempting to evade the innate and adaptive immune response.

## EXPERIMENTAL PROCEDURES

### Cell culture

W6/32 hybridoma cell line (HB-95™, ATCC), A375 cells (CRL-1619™, ATCC) & A375 cells after ERAP1 Knock-Out (KO, previously generated in-house^19^) were cultured in Dulbecco’s Modified Eagle Medium (DMEM, Biowest, L0104) with the addition of 10% Fetal Bovine Serum (FBS, Biowest, S1810), 2 mM L-glutamine (Biowest, X0550) & 1% Penicillin-Streptomycin (Biowest, L022) at 37 °C, 5% CO_2_. The generation of ERAP1 KO clones has been described before^19^. A western-blot analysis confirming the absence of ERAP1 is shown in Figure S1. The immunopeptidome and proteomic analysis of A375 ERAP1 KO cells was performed with clone 1G5. However, later passages of this clone presented gradually unexpected behaviour and morphology. Genome-wide SNP-array analysis^36^ of the 1G5 clone revealed a partial duplication in 1p35 for about 20% of the cells (Figure S2), which however was not reflected in the proteomics results. Subsequently, downstream experiments were performed with clone 1B12 which did not show any genomic instability (Figure S2). THP-1 wild type (TIB-202™, ATCC) and THP-1 ERAP1 KO cells, generated in Dr. Doriana Fruci’s lab at Ospedale Pediatrico Bambino Gesù (Król et al, manuscript in preparation), were cultured in RPMI 1640 (Biowest, L0501) with the addition of 10% FBS (Biowest, S1810), 2 mM L-glutamine (Biowest, X0550) & 1% Penicillin-Streptomycin (Biowest, L022) as usual at 37 °C, 5% CO_2._

### Inhibitor treatment

For the immunopeptidomics, proteomics and surface MHC-I expression analyses, wild type cells were treated either with 10 μΜ ((4-methoxy-3-(N-(2-(piperidin-1-yl)-5-(trifluoromethyl)phenyl)sulfamoyl)benzoic acid), hereafter named as Compound 3^24^, in complete medium (0.1% DMSO) or 0.1% DMSO for 6 days at 37 °C, 5% CO_2_. KO cells (A375 KO:1G5 & THP-1 KO) were cultured in the presence of 0.1% DMSO under the same conditions. During the treatment, the medium was refreshed once. At the end of the treatment, cells were harvested and either subjected to flow cytometric analysis or stored at -80°C until needed for immunopeptidome and proteome isolation. For the rest of the experiments, A375 wild-type and KO (1B12) were treated similarly but the duration of the treatment was reduced to 48 hours.

### Preparation of immunoaffinity columns

The preparation of immunoaffinity columns was performed as previously described ^18,19,37^. Briefly, the W6/32 monoclonal antibody was collected from the HB-95 ™ hybridoma after five days of culture in serum free medium and purified by affinity chromatography using Protein G Sepharose 4 Fast Flow (Cytiva, 17061801). The purified antibody was dialyzed overnight in coupling buffer (NaHCO_3_ 0.1 M, NaCl 0.5 M, pH 8.3). For each column 0.285 g of dry cyanogen bromide activated Sepharose 4B beads (GE Healthcare 17-0430-01) were used. The beads were activated by 1mM HCl, washed with coupling buffer and mixed with 2 mg of purified antibody. The beads with the antibody were rotated overnight at 4°C for coupling. After coupling, beads were washed with coupling and blocking buffer (Tris-HCl 0.1 M, pH 8.0) and blocked for 3h at room temperature. Ultimately, coupled beads were washed with three cycles of acidic (CH_3_COONa 0.1 M, NaCl 0.5 M, pH 4.0) and basic (Tris-HCl 0.1 M, NaCl 0.5 M, pH 8.0) buffer and equilibrated with 20 mM Tris-HCl, pH 7.5, 150 mM NaCl. Pre-columns were prepared in a similar manner with the omission of the W6/32 coupling step.

### Isolation of MHC-I immunopeptidome

The isolation of the MHC-I molecules from A375 cells has been previously described^18,19^. Cell pellets (3-5*10^8^ cells/sample) from wild-type, inhibitor-treated and KO cells (two biological replicates) were thawed on ice and mixed with 20 ml lysis buffer each (Tris–HCl, pH 7.5, 150 mM NaCl, 0.5% IGEPAL CA-630, 0.25% sodium deoxycholate, 1 mM EDTA, pH 8.0, cOmplete™ ULTRA EDTA-free protease inhibitor cocktail tablets (Roche,4134490)) for 1h at 4°C (under rotation). The cell lysate was cleared with ultracentrifugation at 100,000 g for 1 hr at 4 °C and then loaded on the pre-columns and the columns. The flow-through from this procedure was loaded on the columns three more times before washing the columns with 20 bed volumes 20 mM Tris–HCl, pH 8.0, 150 mM NaCl, 20 bed volumes 20 mM Tris–HCl, pH 8.0, 400 mM NaCl, 20 bed volumes 20 mM Tris–HCl, pH 8.0, 150 mM NaCl and finally with 40 bed volumes 20 mM Tris–HCl, pH 8.0. The peptide-MHC-I complexes were eluted by washing with 1% trifluoroacetic acid (TFA).

Elution fractions 1-3 from the above procedure were merged and further purified using reversed-phase C18 disposable spin columns (Pierce™, 89870). Briefly, columns were activated with 50% acetonitrile (ACN) and equilibrated with 5% ACN, 0.1% formic acid (FA). Sample composition was adjusted by adding ACN to a final concentration of 5% and samples were loaded sequentially (150 μl/time) followed by centrifugation (1500 g, 1 minute). Each column was washed with 5% ACN, 0.1%TFA and peptides were eluted by washing two times with 30 μl of elution solution (30% ACN, 0.1% TFA). Due to the presence of β-2 microglobulin in the flowthrough (detected by western blot), the initial eluates were diluted to 5% ACN and the procedure was repeated. The flowthroughs from the C18 column preparations were subjected to Speed-Vac, reconstituted in 5% ACN, 0.1% FA and the C-18 purification was repeated as described above. These samples were analyzed by LC-MS/MS separately.

The purified peptides were further processed by the Sp3 protocol for peptide clean-up ^38^. 20 μg of beads (1:1 mixture of hydrophilic and hydrophobic Sera-Mag™ carboxylate-modified beads, GE Life Sciences) were added to each sample in 95% ACN. Peptide clean-up was performed on a magnetic rack. The beads were washed two times with 100% ACN. Peptides were solubilized in the mobile phase A (0.1% FA in water) and sonicated.

### Isolation of the proteome

Cell pellets were mixed with 150 ml of solution containing 4% sodium dodecyl sulfate, 100 mM Tris/HCl pH 7.6, 0.1M dithiothreitol and incubated at 95◦C for 3 minutes. DNA was sheared by sonication to reduce the viscosity of the sample and then lysate was clarified by centrifugation at 13,000 g for 5 minutes.

The protein extracts were treated with 200 mM iodoacetamide to alkylate reduced cysteine residues and processed according to the Sp3 protocol protocol^38^. 20 μg of beads (1:1 mixture of hydrophilic and hydrophobic SeraMag carboxylate-modified beads; GE Life Sciences) were added to each sample in 50% ethanol. Protein clean-up was performed on a magnetic rack. The beads were washed twice with 80% ethanol followed by one wash with 100% ACN. The beads-captured proteins were digested overnight at 37 ℃ with 0.5 μg trypsin mix in 25 mM ammonium bicarbonate under vigorous shaking (1200 rpm, Eppendorf Thermomixer). The supernatants were collected and the peptides were purified by a modified Sp3 clean-up protocol and finally solubilized in the mobile phase A (0.1% FA in water), and sonicated. Peptide concentration was determined through absorbance measurement at 280 nm using a nanodrop instrument.

### Proteomics Experimental Design and Statistical Rationale

Samples were run on a liquid chromatography-mass spectrometry (LC-MS/MS) setup consisting of a Dionex Ultimate 3000 nano RSLC online with a Thermo Q Exactive HF-X Orbitrap mass spectrometer. Peptidic samples were directly injected and separated on a 25 cm-long analytical C18 column (PepSep, 1.9μm3 beads, 75 µm ID) using an one-hour long run, starting with a gradient of 7% Buffer B (0.1% FA in 80% ACN) to 35% for 40 min and followed by an increase to 45% in 5 min and a second increase to 99% in 0.5min and then kept constant for equilibration for 14.5min. A full MS was acquired in profile mode using a Q Exactive HF-X Hybrid Quadropole-Orbitrap mass spectrometer, operating in the scan range of 375-1400 m/z using 120K resolving power with an AGC of 3x 10^6^ and max IT of 60ms, followed by data independent analysis using 8 Th windows (39 loop counts) with 15K resolving power with an AGC of 3x 10^5^, max IT of 22ms and a normalized collision energy (NCE) of 26.

Immunopeptidomic analysis included two biological replicates per condition (wild-type, inhibitor-treated & ERAP1 KO cells), each analysed 3 times (technical replicates), while a blank sample was also included to increase robustness of peptide identification Immunopeptidomics MS/MS spectra files were searched against the nuORFdb v1.0 database^39^ (323,848 protein sequences) (retrieved:04-03-2024) in Spectronaut® (Biognosys, version 19) using “unspecific” search, with N-terminal acetylation and methionine oxidations as variable modifications and False Discovery Rate (FDR) at 5% at the PSM and peptide level. Spectronaut®’s default settings were chosen for the rest of the parameters (Supplemental Methods). In cases where a peptide was matched to multiple proteins, priority was given to canonical proteins. Comparison between the 3 conditions was performed with an unpaired t-test in Spectronaut®. Two LC-MS/MS runs and searches were performed, as indicated above. Peptides with log2 difference above 1 and with similar behaviour in the two LC-MS/MS runs were considered differentially expressed.

Proteomics analysis included four biological replicates for A375 and three biological replicates for THP-1 cells per experimental condition (i.e. wild-type, inhibitor-treated & ERAP1 KO cells). Raw data were analyzed in DIA-NN 1.8.1 (Data-Independent Acquisition by Neural Networks)^40^ against the reviewed human UniProt protein database (A375 cells: 50516 proteins, retrieved 11-04-2022, THP-1 cells: 20583 proteins, retrieved 8-11-22). Search parameters were set to allow up to two possible trypsin enzyme missed cleavages. A spectra library was generated from the DIA runs and used to reanalyze them. Cysteine carbamidomethylation was set as a fixed modification while N-terminal acetylation and methionine oxidations were set as variable modifications. The match between runs (MBR) feature was used for all the analyses and the output (precursor) was filtered at 0.01 FDR. The protein inference was performed on the gene level using only proteotypic peptides. The double pass mode of the neural network classifier was also activated. Downstream data analysis was performed with the Perseus software v. 1.6.15^41^. Briefly, data were log transformed, identifications with less than 70% valid values in at least one experimental group were removed and missing values were imputed from a normal distribution. For the determination of differentially expressed proteins, Hawaii plots were generated (Pearson correlation, 100 permutations, Class A: S0=0.1, FDR=0.05, Class B: S0=0.5, FDR=0.05). Proteins passing the Class B classification were considered as differentially expressed.

### Isolation of PBMCs

Peripheral blood mononuclear cells (PBMCs) were isolated from a healthy donor according to standard procedures ^42^. Blood was diluted 1:1 with PBS and laid on top of equal amount of Pancoll density gradient medium (Pan Biotech, P04-60500). Samples were centrifuged for 20 minutes at 1200 g without brakes, the buffy coat was collected and washed twice with PBS and cells were either cryopreserved or cultured in 6-well plates at a density of 2,000,000 cells/ml in complete RPMI medium with 100 units/ml interleukin-2.

### Caspase-3/7 assay

For the caspase-3/7 apoptosis assay, A375 cells were seeded at a density of 20,000 cells/ml in a 24-well plate and co-cultured with PBMCs pre-stained with CellTracker Deep Red (Invitrogen) at an effector-target ratio 10:1 for 4 hours in the presence of inhibitor or DMSO. Next, CellEvent™ Caspase-3/7 Green Detection Reagent (Thermo Fisher) was added and apoptotic cells were detected with Leica DMi8 fluorescence microscope (Leica Microsystems) and quantified with ImageJ. Statistical significance was evaluated by one-way ANOVA, followed by Dunnett’s multiple comparisons test in GraphPad Prism v.6.05.

### PBMCs cytotoxicity assay

A375 cells treated with inhibitor were co-cultured with PBMCs pre-stained with CellTracker Deep Red (Invitrogen) at an effector-target ratio 10:1 for 16 hours in the presence of inhibitor or DMSO. After the co-culture, samples were collected, washed and resuspended in 200 μl medium. Prior to acquisition, 50 μl of Count Bright™ Absolute Counting Beads (C36950, Thermo Fisher) along with propidium iodide at a final concentration of 5 μg/ml were added. Data acquisition was performed using a BD LSRFortessa™ flow cytometer (BD Bioscience), and the data were analyzed with FlowJo software (FlowJo, LLC, version 10.10.0)^43^. Statistical significance was evaluated by one-way ANOVA, followed by Dunnett’s multiple comparisons test in GraphPad Prism v.6.05.

### Dichlorofluorescein assay

For the dichlorofluorescein (DCF) assay A375 cells were seeded in black wall, transparent flat-bottom plates at a density of 5,000 cells/well. Cells were exposed to either 0.1% DMSO or Compound 3, as described above. 48 hours later the medium was removed, cells were washed one time with PBS and exposed to either 10 or 25 μΜ DCF and Hoechst 33342 1 μg/ml for 45 minutes at 37 °C, 5% CO_2._ At the end of the incubation period, DCF/Hoechst containing medium was removed, cells were washed twice with PBS and once with Fluorobrite™ DMEM and resuspended in Fluorobrite™ DMEM. Wells exposed to 25 μΜ DCF were used for fluorescence measurement with a BioTek Synergy H1 plate reader (DCF: ex. 485 nm, em. 535 nm/ Hoechst: ex. 352 nm, em. 454 nm). 10 μΜ were used for taking pictures at a Cytation-5 instrument, equipped with filters for YFP (ex. 500, em. 542) and DAPI (ex. 377, em. 477). DCF fluorescence was normalized to Hoechst fluorescence (Synergy H1) or number of nuclei (Cytation 5) and results are expressed as percentage of the wild-type cells.

### Thioflavin-T assay

For the Thioflavin-T (ThT) assay, A375 cells were seeded in a 6-well flat-bottom transparent plate at a density of 75,000 cells/ml (2 ml/well) and treated as described in the caspase assay. At the end of the treatment period, the stock ThT reagent (5mM) was prepared fresh as described in ^44^ in Fluorobrite™ DMEM with 10% FBS, 1% L-glutamine and 1% Sodium-Pyruvate. The medium was then aspirated, cells were washed one time with PBS and were then treated with 5 μΜ ThT in complete Fluorobrite DMEM™ with or without 5 μΜ Dithiothreitol (DTT) for 40 minutes at 37 °C, 5% CO_2._ ThT fluorescence was observed with a Nikon Eclipse Ts2R-FL fluorescence microscope, equipped with a C-LED385* filter (Ex 390/38, BA 475/90). The obtained pictures were pre-processed using Fiji (background subtracted, thresholded). Total intensity was normalized with the area. Results are expressed as percentage of the wild type cells without DTT.

### Mitotracker assay

For the Mitotracker assay A375 cells were seeded in black well, transparent flat-bottom plates at a density of 5,000 cells/well. Cells were exposed to either 0.1% DMSO or Compound 3, as described above. At the end of the treatment, cells were washed one time with PBS and exposed to 200 nM Mitotracker™ Red CMXRos probe (Invitrogen), diluted in Fluorobrite™ DMEM with 1% L-glutamine and 1% Sodium-Pyruvate for 15 minutes at 37 °C, 5% CO_2._ Post-exposure to the probe, the cells were fixed with 4% paraformaldehyde in PBS, stained with DAPI and fluorescence images were taken at a at a Cytation-5 instrument, equipped with filters for Texas Red (ex. 586, em. 647) and DAPI (ex. 377, em. 477). Mitotracker fluorescence was normalized to number of nuclei (Cytation 5) and results are expressed as percentage of the wild-type cells.

### Extracellular Flux Analysis

For the extracellular flux analysis A375 cells were seeded in a Seahorse XF analyzer 24-well plate and incubated for 48 hours with 0.1% DMSO or Compound 3 at 37 °C, 5% CO_2_. Subsequently, cells were incubated for one hour in a non-CO_2_ incubator in DMEM-based assay medium, supplemented with 2mM glutamine and lacking glucose, pyruvate, sodium bicarbonate, HEPES and phenol red. The assay included four injections of 10mM glucose, 1.5μM Oligomycin, 1μM trifluoromethoxyphenylhydrazone (FCCP) and 0.5μM of each of Rotenone and Antimycin A, described in order, and measurements of oxygen consumption rate (OCR) and extracellular acidification rate (ECAR) were collected throughout the assay. Upon completion of the assay, cells were fixed with 4% PFA and stained with DAPI. Normalization was performed by counting nuclei stained with DAPI and results are shown for OCR and ECAR as pmol O_2_/min/cells and mpH/min/cells, respectively.

## RESULTS

### The Immunopeptidome of A375 melanoma cells is responsive to downregulation of ERAP1 activity

To investigate the changes in the immunopeptidome of cancer cells, we compared the peptide repertoire presented by the MHC-I molecules of A375 melanoma cells treated with an allosteric inhibitor targeting the B3P allosteric site of ERAP1 (compound 3 as described in ^24^, also shown in Figure S3) for 6 days at the non-cytotoxic concentration of 10 μΜ (Figure S4) with that of untreated cells and a genetically modified clone lacking ERAP1 expression^19^. Neither the inhibitor treatment nor the ERAP1 KO significantly affected MHC-I cell surface expression as detected by the pan-MHC antibody W6/32 (Figure S5). To isolate the peptide-MHC-I complexes, A375 cells were lysed and the cleared lysate was subjected to immunoaffinity purification with immobilized W6/32 monoclonal antibody. Elution took place under acidic conditions and the purified peptides were analyzed by LC-MS/MS. Due to the relatively low number of peptides identified, we repeated the purification using the flow-through fractions from the first C-18 column and analyzed them in a separate LC-MS/MS run. Since anti-tumor responses can also be targeted against non-canonical peptides, we searched against the Riboseq-based nuORFdb v1.0^39^. In total, 734 unique peptides between 8 and 16 amino acids were identified in all the conditions, after removal of peptides detected in a blank run (Supplemental Table A). Clustering analysis validated the statistical differences between conditions (Figure 1A and Figure S6A). Principle component analysis indicated discreet clusters for each biological condition, suggesting a good degree of reproducibility between the replicates and discrete patterns for each experimental condition (Figure 1B and Figure S6B). Volcano plots comparing the inhibitor-treated A375 cells and ERAP1 KO cells with untreated wild-type A375 cells are shown in Figure 1C-D and Figure S6C-D. These results clearly indicate significant immunopeptidome shifts for both methods of ERAP1 functional disruption.

After merging the two sets of identified peptides, 467 peptides were found to be differentially expressed in the inhibitor-treated cells compared to the control cells (321 peptides significantly upregulated and 146 significantly downregulated, q-value ≤ 0.05 and log_2_fold change ≥1). 501 peptides were differentially expressed in the KO cells compared to the wild-type control (263 peptides significantly upregulated and 238 significantly downregulated). Interestingly, direct comparison between the inhibitor-treated cells and the ERAP1 KO cells indicated that the immunopeptidome shifts were distinct for each method of functional disruption of ERAP1 activity (Figure 1E and Figure S6E). This is likely due to the allosteric nature of the inhibitor and is analogous to previous results obtained with an ERAP1 inhibitor targeting the malate allosteric site^19^. This finding reinforces the notion that targeting distinct allosteric sites in ERAP1 may be a viable method for fine-tuning the immunopeptidome for therapeutic applications.

**Figure 1:**
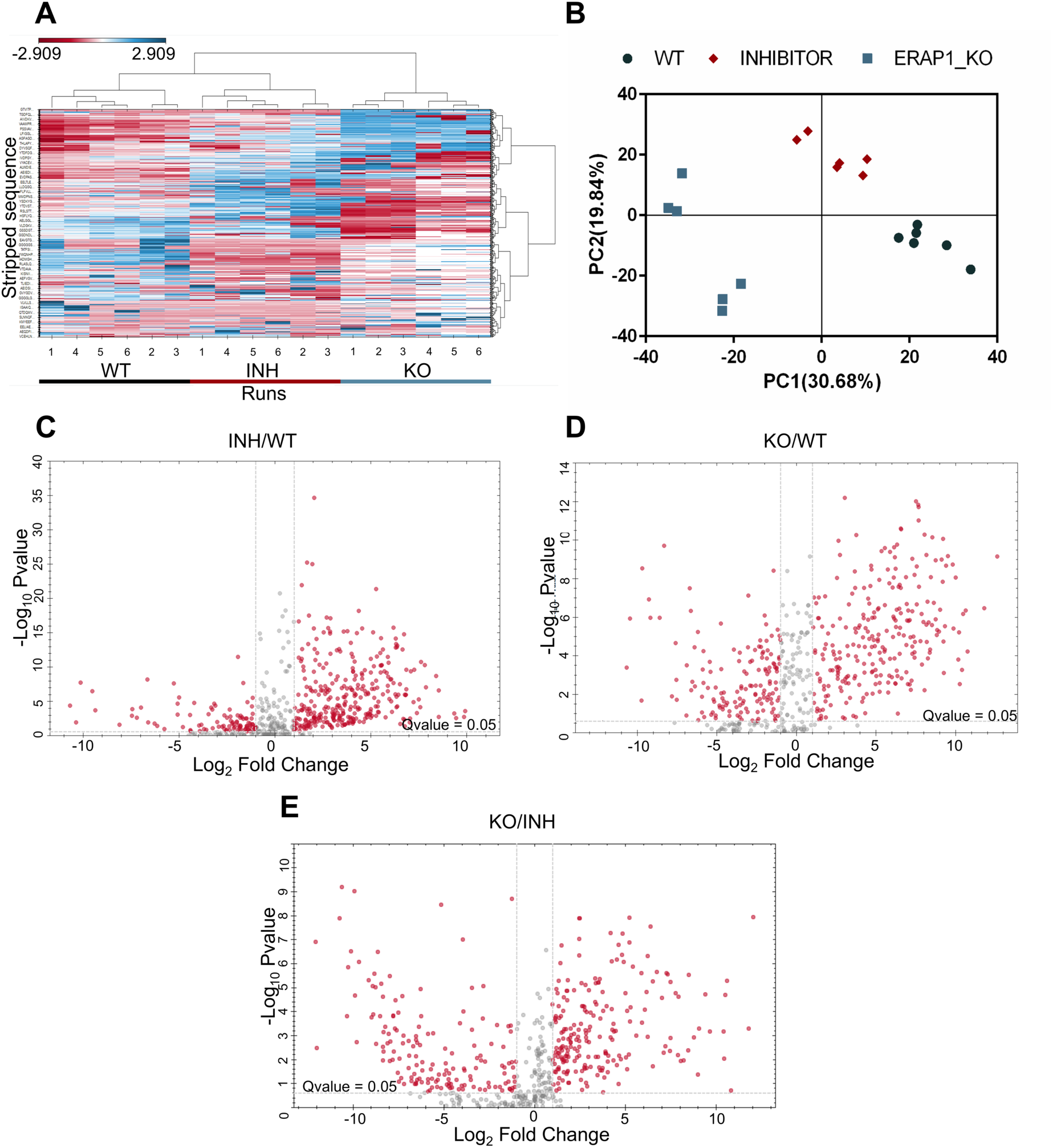
Analysis of the immunopeptidome of wild-type, inhibitor-treated & ERAP1 KO A375 cells. **Panel A**, Heatmap from LC-MS/MS run 1 showcasing peptide distribution across different experimental conditions. Hierarchical clustering was calculated by Manhattan Distance. Colors indicate peptide intensities, ranging from low (red) to high (blue). (Graph generated with Spectronaut® v. 19). **Panel B,** Principal Component analysis (PCA) from LC-MS/MS run 1. PCs 1 & 2 contribute to the explanation of 48% of the sample variability. Based on these two PCs, wild-type samples (circle), inhibitor-treated samples (diamond) and KO samples (square) formed 3 distinct groups, indicating that the replicates within each group share similarities with each other but there were differences in the immunopeptidomes between different treatments. **Panels C-E,** Volcano plots from LC-MS/MS run 1 indicating the statistical significance of the differences between (C) the inhibitor-treated and the wild type A375 cells, (D) the genetically modified (KO) and the wild type A375 cells and (E) the KO versus inhibitor-treated cells. Each circle represents a unique peptide sequence. Peptides with a q-value ≤0.05 and a log2 fold change ≥1 are considered statistically significant.

To estimate the binding affinity of the identified peptides to the HLA-I alleles expressed by the A375 cells (HLA-A*01:01, A*02:01, B*44:03, B*57:01, C*06:02, C*16:01) we used the HLAthena affinity prediction server for peptides between 8 and 11 amino acids long ^45^. Figure 2A shows that most of the peptides are predicted to bind to the HLA-I alleles carried by A375 cells, as opposed to a random set of peptides that was used as a control for the predictions. No major differences were observed between conditions, suggesting that ERAP1 functional disruption does not grossly block antigen presentation. However, a substantial shift towards longer-length peptides was evident (Figure 2B) in the inhibitor-treated and ERAP1 KO, consistent with the inhibition of the peptide-shortening activity of ERAP1. Interestingly, the allosteric inhibitor targeting the B3P site used here affected peptide length almost as much as the ERAP1 KO, in contrast to the effect of a previously analyzed allosteric inhibitor targeting the malate allosteric site^19^. This finding suggests that different allosteric sites in the internal cavity of ERAP1 may play distinct roles in length selection^46^.

To better evaluate whether the observed immunopeptidome shifts may translate to changes in the immunogenicity of cells, we predicted the immunogenicity score of the differentially expressed 9mers using the DeepImmuno immunogenicity prediction tool^47^. The distribution of the obtained immunogenicity scores indicated that a higher percentage of the presented 9mers upregulated in the KO and inhibitor-treated cells had scores above 0.8, when compared to the downregulated peptides (Figure S7). Moreover, we searched the identified peptides to see if they correspond to known antigenic peptides listed in the Internet epitope database (IEDB). We identified several known antigenic peptides (including MAGE antigens and antigenic epitopes listed in the IEDB), several of which were upregulated by ERAP1 KO or inhibition (Supplemental Tables S1 and S2). Interestingly, several antigenic peptides were altered in non-identical ways between KO or inhibitor-treated cells suggesting that for these antigenic epitopes the two ways of functional disruption may not be equivalent. Interestingly, 2.3% of the identified peptides originated from unannotated proteins (Supplemental Table S3), some of which were upregulated by ERAP1 KO or inhibition. These results suggest that blocking the processing of antigenic peptides by ERAP1 can lead to specific immunopeptidome changes that can promote cellular antigenic responses and this can include unannotated protein ORFs that can be a source of neoantigenic epitopes.

**Figure 2.**
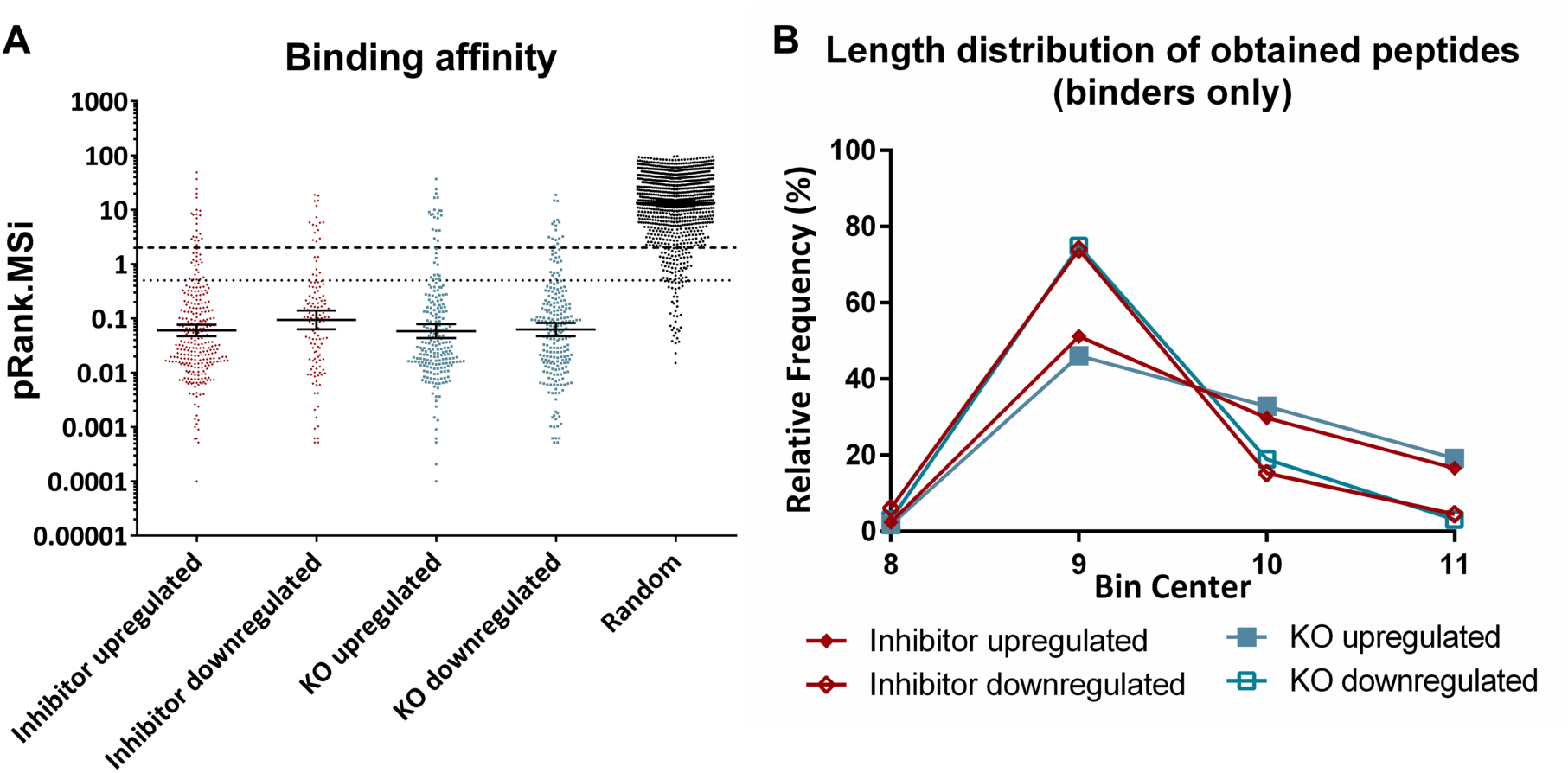
**Panel A**, Distribution of predicted affinities (HLAthena) of the significantly up-or downregulated peptides after inhibitor treatment or ERAP1 KO for the HLA alleles present in A375 cells (A*01:01, A*02:01, B*44:03, B*57:01, C*06:02, C*16:01). Each point on the graph signifies a distinct peptide sequence. Only the score for the top predicted HLA allele is depicted for each peptide. A set of random peptide sequences generated by RandSeq (https://web.expasy.org/randseq/) was also plotted as negative control. Prediction scores below 2 (dashed line) indicate binding to at least one of the HLA alleles, with scores below 0.5 (dotted line) indicating strong binding. **Panel B,** Length distribution of the significantly up- or downregulated binding peptides after inhibitor treatment or ERAP1 KO.

### ERAP1 functional disruption can enhance the immunogenicity of tumor cells

Immunopeptidome shifts due to ERAP1 inhibition have been theorized to affect the recognition of cancer cells by CD8+ T cells by either enhancing the presentation of rare antigenic epitopes^48^, by inducing the presentation of novel epitopes that elicit the generation of new effector T cells^49^, or by perturbing the interaction with inhibitory NK receptors ^50–52^. To test whether the immunopeptidome shifts described above can affect cytotoxic responses against this melanoma cell line, we co-cultured wild-type A375 cells, ERAP1 KO A375 cells or A375 cells treated with the allosteric inhibitor with human PBMCs. Cytotoxic responses of immune cells were evaluated by a fluorescence, microscopy-based, caspase apoptosis imaging assay and a flow cytometry-based cytotoxicity assay. Compared to wild-type cells, both assays showed a similar trend of increased killing of inhibitor-treated and ERAP1 KO cells (Figure 3). Although the cytotoxicity enhancement was mild, it was consistent throughout both methods of ERAP1 functional disruption and both functional assays. The magnitude of functional responses observed was limited, especially compared to previous *in vivo* studies^48,50^. However, it should be noted that this assay can only report on the enhancement of pre-existing or cross-reacting cellular responses, as the amplification of T-cell clones is not possible, thus minimizing potential effects. Overall, this result reinforces our confidence that blocking ERAP1 function can, even in the absence of combinatorial treatments, contribute to enhanced immunogenicity.

**Figure 3.**
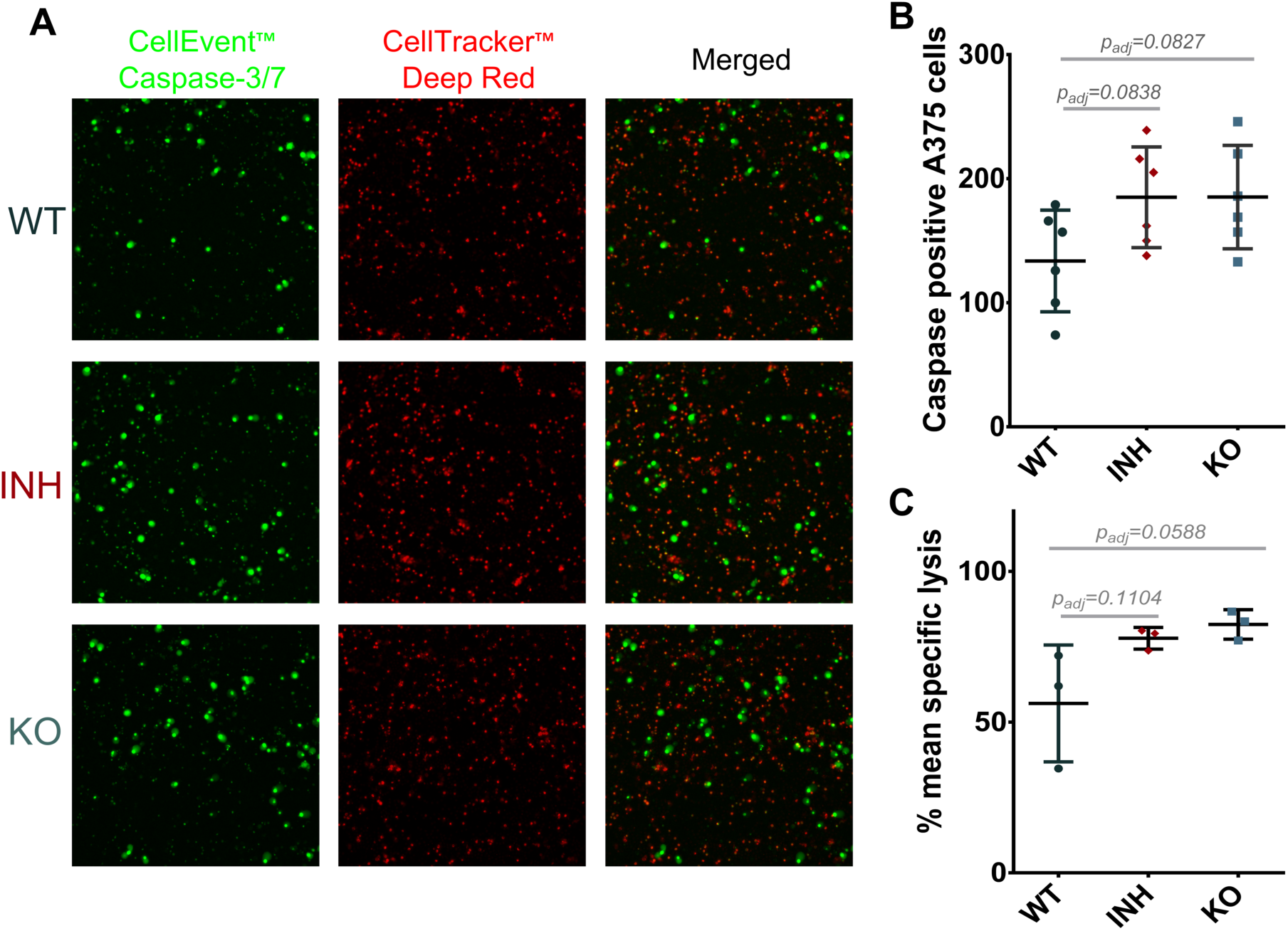
**Panel A**, representative fluorescence microscopy images of activated PBMCs (red) and caspase positive A375 cells (green) in all conditions. **Panel B,** Graph showing the changes in caspase-positive inhibitor-treated (diamond) and ERAP1 KO (square) A375 cells, compared to A375 wild-type cells (circle). The average, with its respective standard deviation, was calculated from six images per experimental condition. Statistical significance was evaluated by one-way ANOVA, followed by Dunnett’s multiple comparisons test in GraphPad Prism v.6.05. **Panel C,** Graph showing the change in mean-specific lysis in inhibitor-treated and ERAP1 KO cells in the PBMC cytotoxicity assay. The average, with its respective standard deviation, was calculated from three technical replicates per experimental condition. Statistical significance was evaluated by one-way ANOVA, followed by Dunnett’s multiple comparisons test in GraphPad Prism v.6.05.

### Functional disruption of ERAP1 alters the proteome of A375 cells

To examine if ERAP1 activity has significant effects on the proteome of the A375 cell line, we used data-independent analysis to compare the proteomic content of the same conditions we used for the immunopeptidome analysis described above. In total, 12 replicates per experimental condition were analyzed (4 biological replicates, 3 technical replicates for each biological). Overall, we identified 5575 proteins, 5049 of which remained after filtering (Supplemental Table B). 1375 of those proteins were differentially expressed in the inhibitor-treated and ERAP1 KO cells (FDR= 0.05, S0= 0.5). Based on the identified proteins, the 3 conditions tested (WT, inhibitor-treated and KO cells) formed discreet hierarchical clusters (Figure 4A) and grouped in the principal component analysis (Figure 4B), which suggests distinctive features for each experimental condition. 494 proteins were differentially expressed in the inhibitor-treated cells compared to wild-type cells (Figure 4C). 74% (367 out of 494) of those proteins were also significantly altered similarly in the KO cells. 1252 proteins were differentially expressed in the KO cells compared to wild-type cells (Figure 4D). Although both methods of ERAP1 functional disruption resulted in significant proteome shifts, they were not identical. Indeed, more proteins were both upregulated and downregulated in the KO compared to the inhibitor-treated cells (Figure S8). This could either indicate non-complete inhibition of ERAP1 by the compound, or indirect effects on the proteome due to the lack of the protein in the KO cells. Several components of the antigen presentation pathway were found altered, primarily in the ERAP1 KO cells, such as components of the PLC (PDIA3, TAPBP, CALR and TAP2), suggesting potential compensatory changes in the antigen loading process due to lack of ERAP1 (Figure 4C-D). Interestingly, we detected changes in abundance for the MHC alleles HLA-A and HLA-DRB3, which may reflect changes in the peptide repertoire available for their cargo. In addition to these, we observed alterations in other related processes, such as protein folding, where several proteins were upregulated (HSP90AA1, PPIA in both inhibitor and KO; HSPA4L, FKBP10 and SIL1 only in the KO, Figure 4E), and proteasomal degradation, where several proteins were downregulated (PSMB2, PSMD6, PSMD10, UBE2A, PCBP2, SKP1, BAG6, UBQLN1 in both the inhibitor and KO; PSMD14, PSME1, RAD23A, UBQLN2 only in the KO, Figure 4F). Interestingly, proteins related to interferon signaling were also affected by ERAP1 inhibition (Figure 4G).

**Figure 4.**
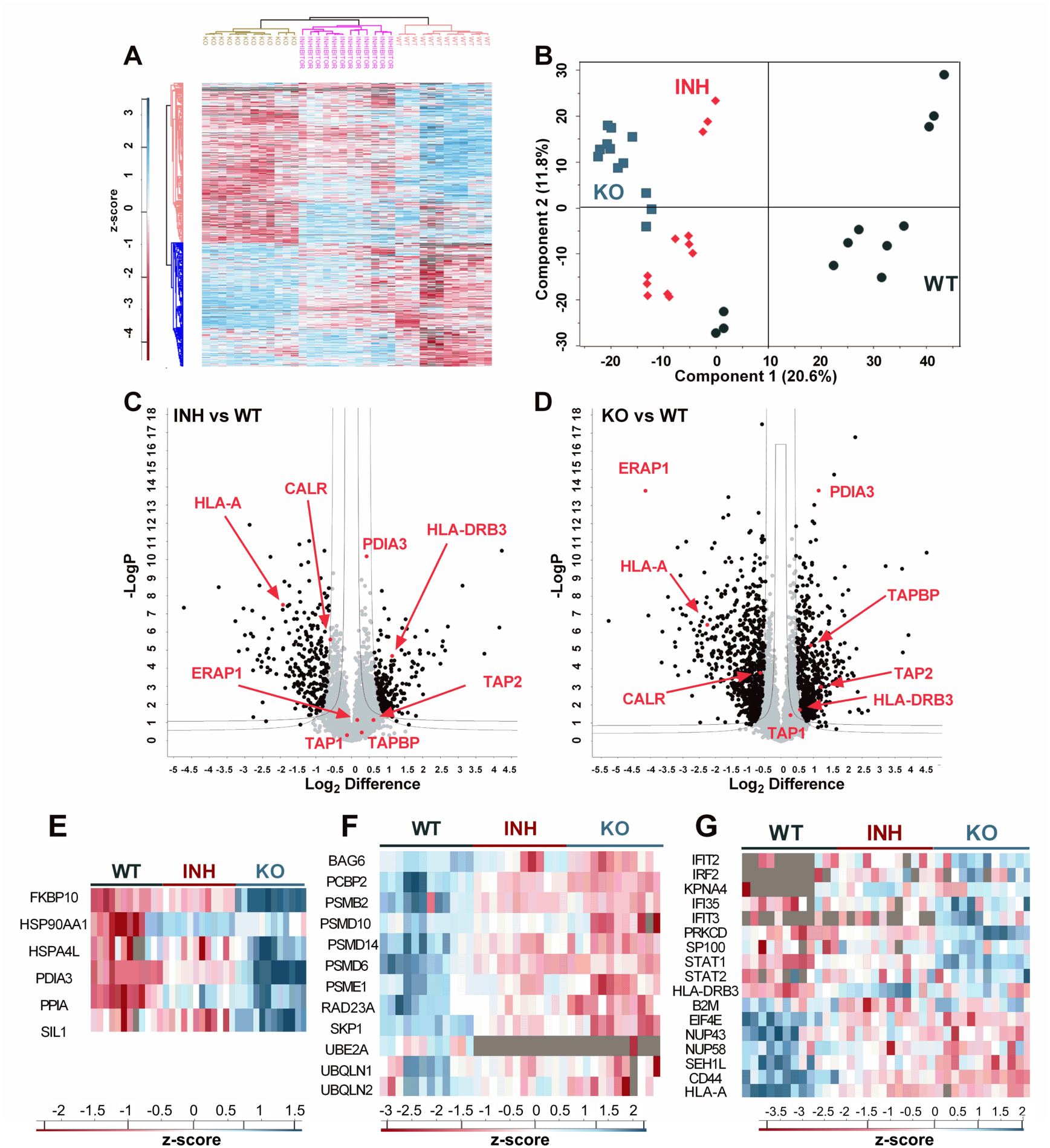
Proteomic analysis of wild-type, inhibitor-treated & ERAP1 KO A375 cells. **Panel A**, Heatmap showing the cluster analysis and distribution of proteins in the three conditions (WT, inhibitor treated & ERAP1 KO A375 cells) for each replicate (four biological replicates, each measured in three technical replicates) (Graph generated with Perseus v. 1.6.15). **Panel B**, Principal Component Analysis (PCA) of the three experimental conditions. Each point represents a distinct biological or technical replicate. **Panels C & D**, Hawaii plots of identified proteins, indicating the statistical significance of the observed differences in protein abundance between the two treatment conditions (inhibitor-treated & ERAP1 KO) and the WT cells. 494 proteins in the inhibitor-treated (C) and 1252 proteins in the KO cells (D) were differentially expressed. Select proteins that participate in antigen presentation are indicated in red. (Graph generated with Perseus v. 1.6.15) **Panels E, F & G**: Heatmaps of proteins participating in protein folding (E), proteasomal degradation (F) and interferon signaling (G).

Changes in the proteome may also affect the composition of the immunopeptidome. Although it is now well established that ERAP1 can regulate the cellular immunopeptidome through its ER trimming activity, its effects on the cellular proteome may also contribute indirectly. Indeed, of the 501 peptides found to be differentially presented in the ERAP1 KO cells, 79 (15.8%) of them belong to proteins that have altered expression levels in the same cells. This suggests that proteomic changes due to ERAP1 KO can affect up to 1/6 of the immunopeptidome. In terms of proteins however, of the 1252 proteins that were found to be expressed in different levels in the ERAP1 KO cells, only 63 (5.0 %) were associated with differentially presented peptides in the immunopeptidome, indicating that the impact of the proteomic changes on the immunopeptidome is limited to a small subset of cellular proteins. These effects were smaller in the case of the ERAP1 inhibitor (6.9% and 4.9% respectively). These findings suggest that although ERAP1 activity can influence the cellular proteome and some of those changes can be translated to immunopeptidome shifts, they are likely secondary to the direct effect of ERAP1 on the generation or destruction of antigenic peptides in the ER.

Given the observed proteomic changes upon ERAP1 disruption, we explored if those changes translate to specific biochemical pathways. We used the ShinyGΟ tool ^53^ to compare between the two treatment conditions. Differentially expressed proteins in each experimental condition were used as input to identify relevant KEGG ^54^ and Reactome ^55^ pathways. The top 20 pathways ordered by fold-enrichment are shown in Figure S9. Overall, despite the existence of some unique pathways for each experimental condition, several of the pathways are shared between the inhibitor-treated and KO cells. These include, among others, oxidative phosphorylation, chemical carcinogenesis-reactive oxygen species, metabolic pathways and several disease-related pathways for the KEGG database. Cellular responses to stress and several transport-related pathways were also identified by the Reactome database.

### ERAP1 disruption affects the proteome of THP-1 cells

Given the surprisingly large effect of both genetic and pharmacological inhibition of ERAP1 on the proteome of A375 cells, we inquired whether this phenomenon is specific to A375 cells or is more general to other cancer cells. To address this, we analyzed the proteomic changes of THP-1 cells, treated identically to A375. For THP-1 cells we identified 5758 proteins (5391 after filtering, Supplemental Table B). 1594 were differentially expressed in the inhibitor-treated or KO cells, compared to the WT control (FDR= 0.05, S0=0.5). Similar to the results obtained for A375 cells, hierarchical clustering and principal component analyses indicated three distinct clusters consistent with distinct proteomic profiles per biological condition (Figure S10 A and S10 B). 209 proteins were differentially expressed in the inhibitor-treated cells and 158 of them were affected in a similar manner in the ERAP1 KO. ERAP1 KO altered the levels of 1548 proteins (Figure S10 C and S10 D). Venn diagrams indicate that despite the smaller effect of the inhibitor when compared to the KO cells, there are several shared changes in the significantly up-regulated and down-regulated proteins between the two treatment conditions (Figure S10 E and S10 F).

By comparing the proteomic changes observed in the A375 and THP-1 cell lines, we identified common patterns in both cell lines. In particular, some components of the PLC were similarly affected. For example, the Transporter Associated with Antigen Processing chain 2 (TAP2), a component of the TAP complex responsible for the transport of peptides from the cytosol to the endoplasmic reticulum ^56^, was found upregulated in both A375 and THP-1 ERAP1 KO cells (Figure 4D and Figure S10D). Tapasin (TAPBP) and calreticulin (CALR) followed a similar trend. Furthermore, in both cellular systems ERAP1 KO led to significant reductions in the levels of some HLA (HLA-A on A375 cells and HLA-B on THP-1 cells). Pathway analysis of the proteomic changes in THP-1 cells (Figure S11) suggested several common pathways with A375 cells, including oxidative phosphorylation, chemical carcinogenesis-reactive oxygen species, metabolic pathways, cellular response to stress and transport related pathways. Taken together, our analyses suggest that the effects on the proteome of cancer cells induced by ERAP1 inhibition or ERAP1 KO are not limited to the A375 cell line but may extend to other cancer cell types as well.

### ERAP1 inhibition reduces reactive oxygen species in A375 cells

Since the proteomics results and the subsequent pathway analysis (Figure S9 & S12) suggested that ERAP1 function may relate to changes in cellular stress and reactive oxygen species (ROS), we used the dichlorodihydrofluorescein (DCF) dye to assess the activity of ROS in wild-type A375 cells in comparison to ERAP1 KO and inhibitor treated cells (Figure 5). Wild-type A375 cells showed a significant DCF signal indicating substantial ROS activity, a common feature of cancer cells^57^. Surprisingly, blocking ERAP1 function either by inhibitor or by genetic silencing resulted to a substantial decrease in DCF signal, indicating a reduction of ROS activity when ERAP1 activity is blocked (Figure 5 B and 5 C). This finding suggests that in A375 cancer cells, ERAP1 activity may promote ROS formation.

**Figure 5:**
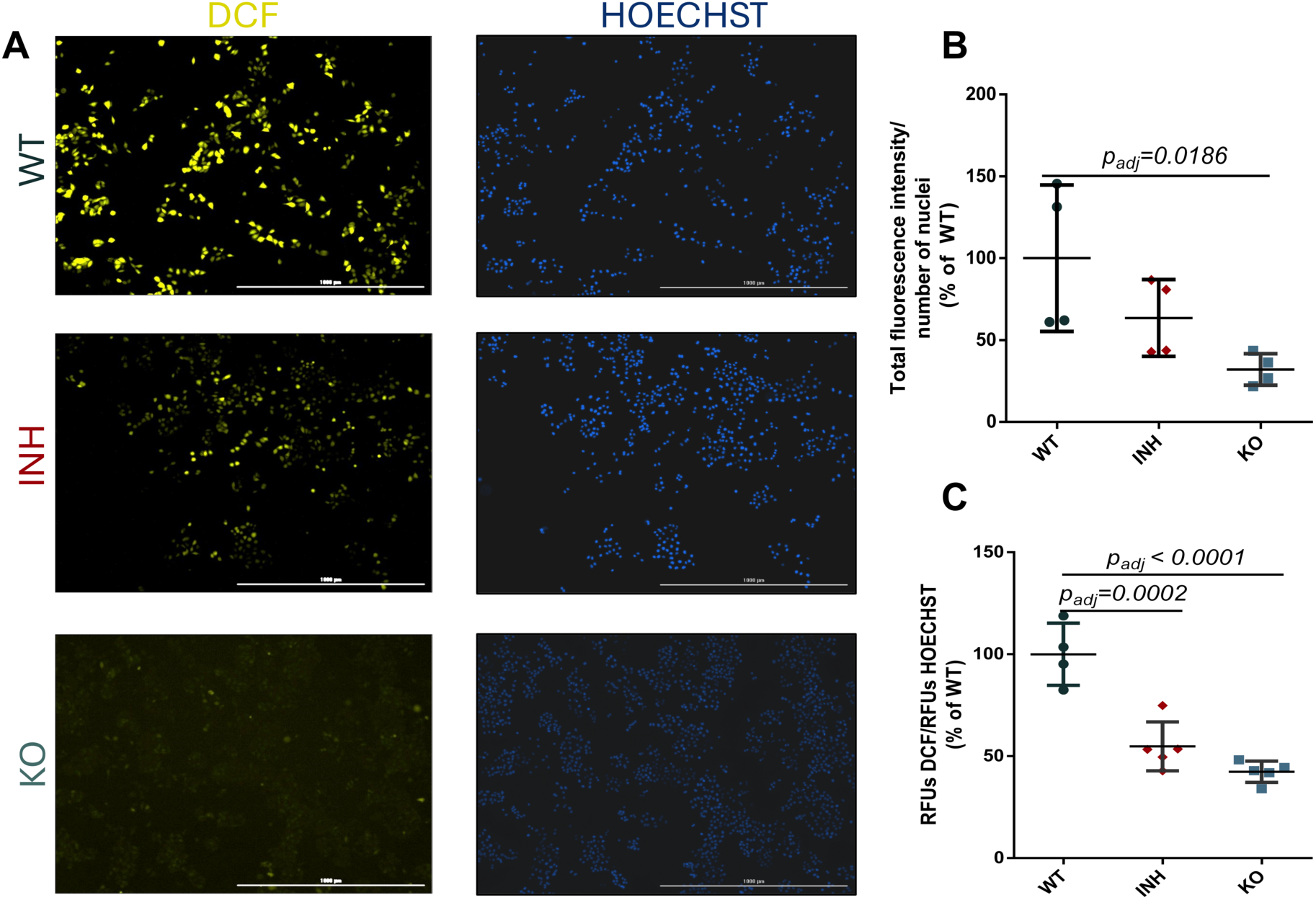
Dichlorodihydrofluorescein (DCF) assay for measurement of reactive oxygen species (ROS). **Panel A**, representative images of wild-type (WT), inhibitor-treated (INH) and ERAP1 KO (KO) A375 cells incubated with DCF (left) or Hoechst 33342 (right). **Panel B,** total fluorescence intensity per number of nuclei quantitated by direct cell imaging on a Cytation-5 instrument for the three biological conditions. **Panel C,** relative DCF fluorescence units (RFU) normalized to Hoechst dye fluorescence measured by direct fluorescence measurement on a BioTek Synergy H1 plate reader. Statistical significance was evaluated by one-way ANOVA, followed by Dunnett’s multiple comparisons test in GraphPad Prism v.6.05.

### ERAP1 inhibition sensitizes cells to external ER stress

The observations of the pathway analysis combined with the effect of ERAP1 inhibition on ROS formation, prompted us to investigate the potential effects of ERAP1 activity in ER stress. Indeed, ERAP1 activity in the ER has been associated with ER stress in the case of the MHC-I allele HLA-B*27, leading to misfolding and regulating the generation of HLA-B*27 allele heavy-chain dimers, which induce autoreactive-like immune responses^58–60^. However, due to the unusual folding properties of HLA-B*27, it is not known if similar effects can occur in different cellular contexts. To investigate this, we used the Thioflavin T (ThT) assay. ThT exhibits increased fluorescence upon binding to protein aggregates and can thus serve as an indicator of ER protein misfolding and ER stress^44^. To evaluate the capability of the cells to compensate for external stressors we used DTT, a compound that can interfere with the redox potential, disulfide bond formation and protein folding in the ER. The ThT fluorescence was similar between wild-type, ERAP1-KO and inhibitor-treated cells (Figure 6). The addition of DTT, however, led to a significant increase in the signal from wild-type cells, which was much more pronounced in inhibitor-treated and ERAP1-KO cells, indicating that the inhibition of ERAP1 activity, lead to a sensitization of the cells to an external ER stressor.

**Figure 6:**
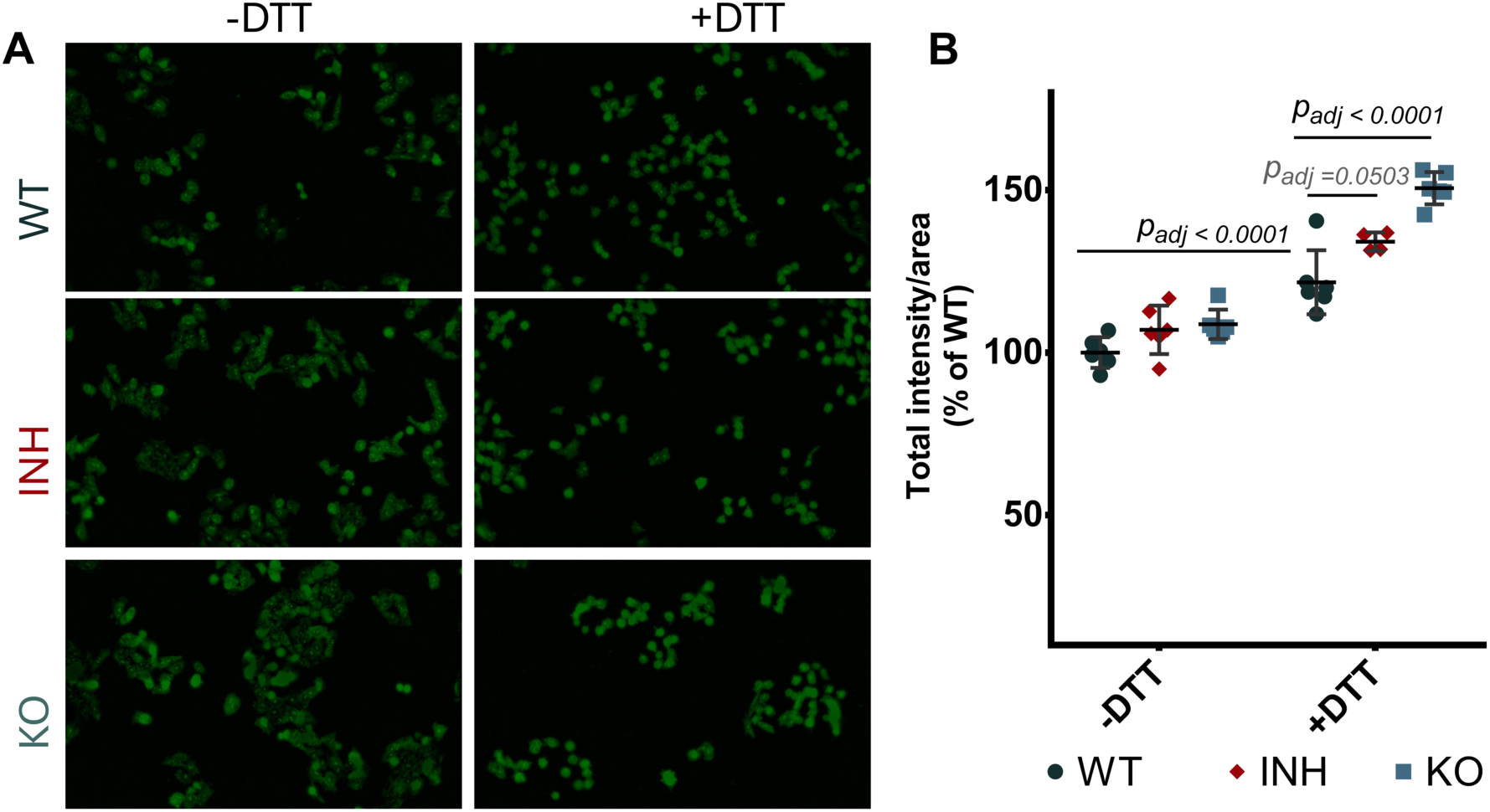
Thioflavin T assay for ER stress. **Panel A**, representative images of wild-type (WT), inhibitor-treated (INH) and ERAP1 KO (KO) A375 cells incubated with Thioflavin T after treatment with DTT as described in the Experimental Methods section. **Panel B,** total fluorescence intensity normalized for cell area for the three biological conditions in the absence or presence of DTT. Statistical significance was evaluated by one-way ANOVA, followed by Dunnett’s multiple comparisons test in GraphPad Prism v.6.05.

### ERAP1 can indirectly affect metabolism and mitochondrial function

Inspired by the pathway analysis which suggested potential changes in oxidative phosphorylation pathways, we identified several relevant proteins in our dataset (Figure S13) and we used the Mitotracker™ Red CMXRos probe to explore potential changes in mitochondrial potential^61^. Mitochondria can communicate and cooperate with the ER to maintain cellular homeostasis^62^. We incubated ERAP1 KO and inhibitor-treated A375 cells with the Mitotracker™ probe before fixation and DAPI staining and normalized the probe’s fluorescence with the number of nuclei. (Figure 7 A). Interestingly, both ERAP1 inhibition and ERAP1 KO, resulted in a significant decrease of Mitotracker signal (Figure 7 B) suggesting a reduction in mitochondrial membrane potential (Δψ_μ_) upon ERAP1 functional disruption. This finding could indicate that loss of ERAP1 activity in the ER indirectly affects mitochondrial function leading to partial depolarization, which can then affect oxidative phosphorylation.

**Figure 7:**
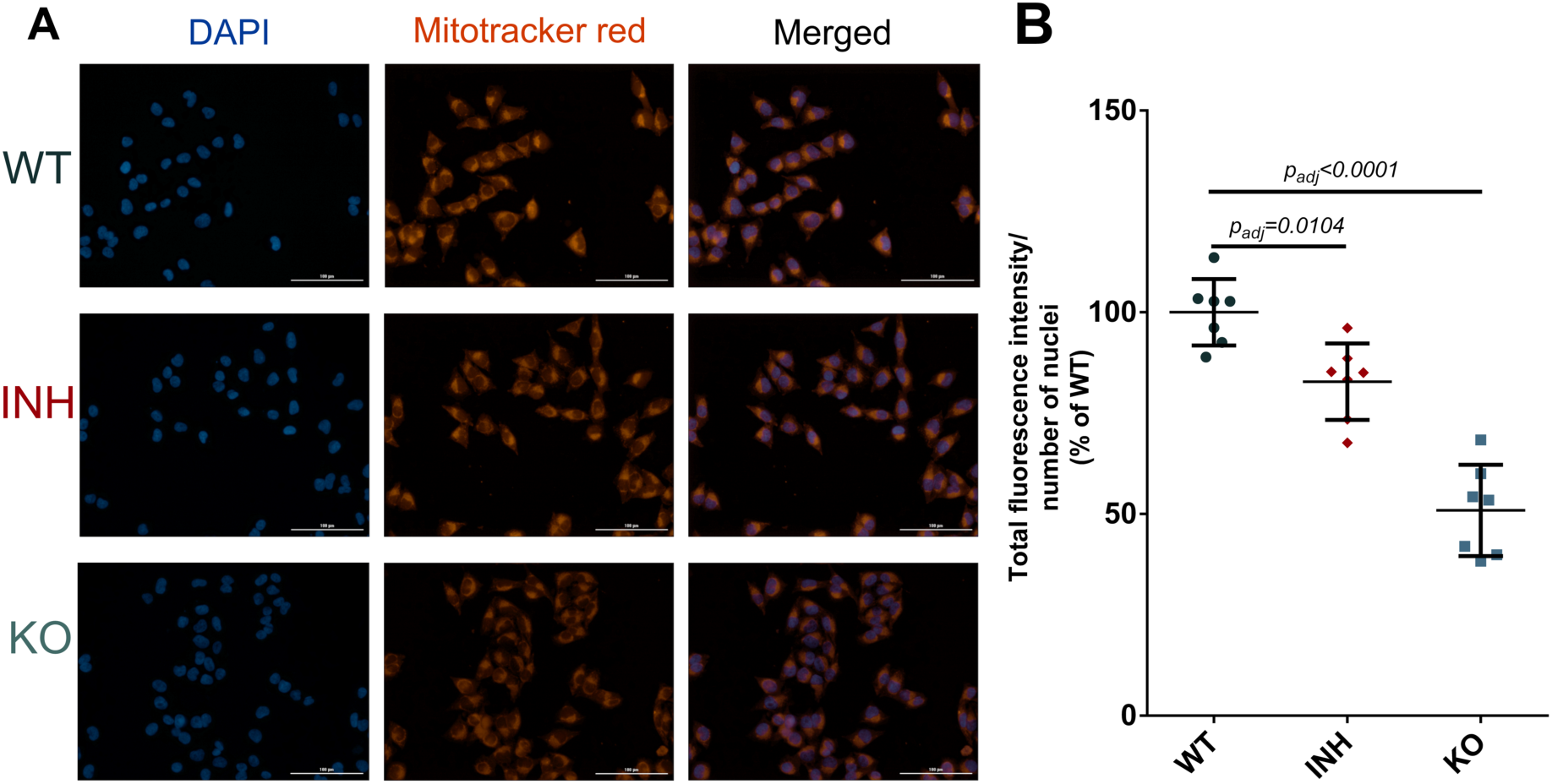
**Panel A**, representative fluorescence microscopy images of A375 cells (wild-type, inhibitor treated or ERAP1 KO) visualized after staining with DAPI (blue) and Mitotracker (red) as well as merged. **Panel B,** mean fluorescence of Mitotracker per identified nuclei for the three conditions. The calculated adjusted p values comparing KO cells and inhibitor treated cells to the wild-type cells are indicated. Statistical significance was evaluated by one-way ANOVA, followed by Dunnett’s multiple comparisons test in GraphPad Prism v.6.05.

### Extracellular flux analysis

To better understand the potential impact of ERAP1 function on mitochondrial metabolism we used the Seahorse XF analyzer to measure Oxygen Consumption Rate (OCR) and Extracellular Acidification Rate (ECAR) in wild-type (WT), inhibitor-treated (INH), and KO cells. This experiment can evaluate mitochondrial respiration and glycolytic activity under different conditions and metabolic stressors, revealing differences in the mitochondrial metabolic profile when ERAP1 function is disrupted (Figure 8). When analyzing the OCR profile, while several parameters were not affected (Figure 8, panels C-E), we observed a statistically significant increase in the basal respiration levels (Figure 8B), proton leak (Figure 8F) and non-mitochondrial respiration (Figure 8G) in the ERAP1 KO cells. A trend towards increased non-mitochondrial respiration was also observed in the inhibitor-treated cells, although much less pronounced. This finding may suggest some limited mitochondrial dysfunction which is in line with changes in Mitotracker signal described above. Alternatively, this finding may indicate increases in energy demand that cannot be easily satisfied by regular mitochondrial respiration. Accordingly, our analysis revealed some increases in basal glycolytic levels, spare and maximum glycolytic capacity (Figure 8, panels I-K) also indicating small changes in energy production in the ERAP1 KO cells. Taken together, these changes suggest that loss of ERAP1 may induce some metabolic pressure onto mitochondria that leads to proteomic and metabolic responses.

**Figure 8:**
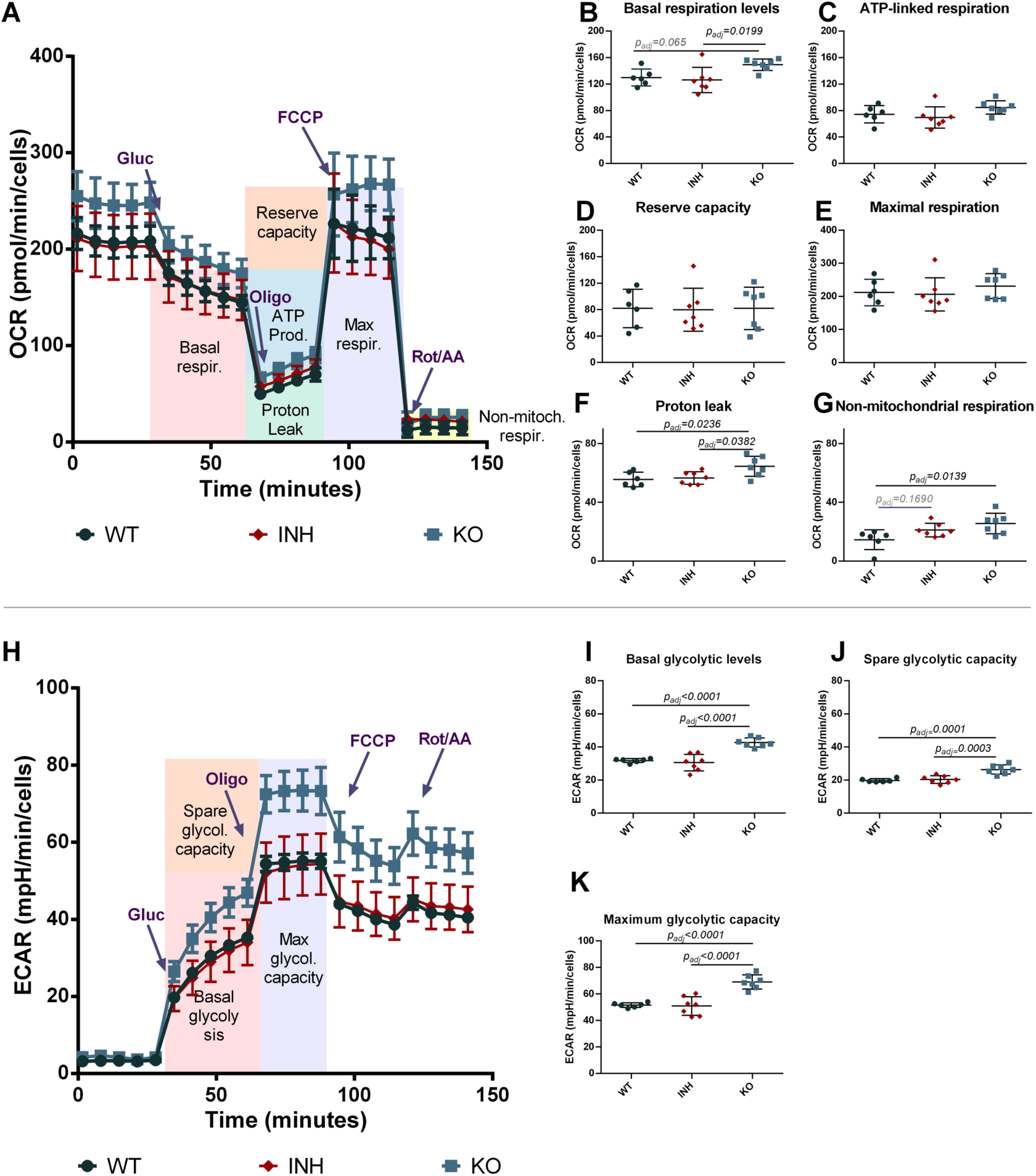
Seahorse™ extracellular flux analysis of effects of ERAP1 disruption on cellular metabolism. **Panel A**, Oxygen Consumption Rate (OCR) followed as a function of time during the experiment for wild-type, inhibitor-treated and ERAP1 KO A375 cells. Arrows at specific time points indicate the addition of Glucose, Oligomycin, Carbonyl cyanide-p-trifluoromethoxyphenylhydrazone (FCCP) and Rotenone / Antimycin A to probe different metabolic components, as indicated by the colored regions in the figure. **Panels B-G**, quantification and comparison of components of mitochondrial metabolism between the three experimental conditions. **Panel H,** Extracellular Acidification Rate (ECAR) followed as a function of time as in panel A. **Panels I-K,** quantification and comparison of basal glycolytic levels, spare glycolytic and maximal glycolytic capacity as calculated form the ECAR measurements. Statistical significance was evaluated by one-way ANOVA, followed by Dunnett’s multiple comparisons test in GraphPad Prism v.6.05.

## DISCUSSION

ERAP1 has been primarily studied for its role in generating antigenic peptides and is considered to have a significant specialization for this role. However other roles have been proposed, albeit less well characterized^25–33^. Indeed, metallo-aminopeptidases such as ERAP1, often have multiple biological functions depending on the cellular context, tissue, cell type, cellular compartment, and pathophysiological context^63^.

Here, we combined immunopeptidomics with proteomic, biochemical, and metabolic analyses to better understand the effect of ERAP1 activity on cancer cells. We find that, consistent with previous reports, ERAP1 is a key regulator of the cellular immunopeptidome and the mechanism of ERAP1 modulation is reflected on the nature of the immunopeptidome. We furthermore show that regulation of the immunopeptidome results to modest increases in immunogenicity, which could be a consequence of alterations in the levels of antigenic peptides and potential neoantigens originating from either annotated or unannotated protein ORFs.

To our surprise, we also find that ERAP1 inhibition induces significant shifts in the cellular proteome. Since the immunopeptidome is generated from the sampling of cellular proteins, proteome shifts may contribute to immunopeptidome shifts. However, the extent to which the immunopeptidome represents the cellular proteome is a matter of significant debate. Several studies have argued on whether the immunopeptidome is a real-time representation of all proteins in the cell or a directed sampling of the dynamic changes in protein synthesis^8,64^. Here, we find that ERAP1 genetic silencing or pharmacological inhibition has repercussions on the cellular proteome and that these proteomic changes contribute to the immunopeptidome changes only to a limited extent. Less than 1 out of 6 peptides in the immunopeptidome belong to proteins that are altered by ERAP1 inhibition and less than 1 out of 20 proteins that are affected by ERAP1 inhibition carry peptide sequences that are altered in the immunopeptidome. This discrepancy between the proteome and immunopeptidome shifts suggests that some proteins may be preferentially represented in the immunopeptidome, which is consistent with the idea that antigen presentation is not an unbiased constant sampling of the cell proteome but rather a more directed process, reporting select changes in protein content upon infection^8^. Spatial compartmentalization of antigen presentation may also contribute to this specialization^65^. Moreover, the limited contribution of the proteome to the immunopeptidome indicates that the effects of ERAP1 on antigen presentation are dominated by its direct role in preparing antigenic peptides over indirect effects on the cellular proteome. In this context, these results strengthen the notion that ERAP1 is a specialized enzyme in editing the immunopeptidome.

Despite the small representation of proteomic changes on the cell surface, the scale of some of these changes was surprisingly large, indicating a secondary, but potentially important, role of ERAP1 on cellular homeostasis. This effect was also reproduced to a reasonable extent in another cancer cell line, suggesting that this is not an isolated property of A375 cells but could be a fundamental consequence of the enzyme’s function in maintaining peptide homeostasis in the ER, which could affect protein folding and indirectly contribute to ER stress. Indeed, in the case of the MHCI-allele dependent inflammatory disease, Ankylosing Spondylitis, the pathogenetic role of ERAP1 has also been proposed to be linked to ER misfolding of HLA-B*27, the formation of heavy chain dimers and general ER stress^59,60^.

It should be noted, however, that the cellular systems explored here (A375 and THP-1 cells) are both cancer cell lines which may not represent normal cell metabolism. Although our observations may not extend to non-cancer cell lines, ERAP1 is a pharmacological target in cancer therapy and any effects of its inhibition on cancer cell function may have to be validated and evaluated separately to its effect on the immune responses. For this reason, we built on proteomic findings with additional biochemical and metabolic analyses. We find that in the A375 melanoma cell line, ERAP1 functional disruption appears to affect the capability of the ER to respond to redox challenges. These effects are accompanied by changes in the levels of reactive oxygen species and metabolic changes focused on mitochondrial membrane polarization and extending to glycolytic potential and non-mitochondrial respiration. Specifically, we observed a mild increase in proton leak in the ERAP1 KO cells, in addition to a decrease in mitochondrial membrane potential (Δψ_μ_) and reactive oxygen species in both treatment conditions. Mild proton leak has been suggested as a protective mechanism against oxidative damage by reducing ROS formation^66–69^ due to a lower mitochondrial membrane potential (Δψ_μ_) ^69^. This mechanism may prevent excessive electron supply to the respiratory chain, thereby lowering the chance of electron leakage and subsequent superoxide production. Moreover, a trend towards increased non-mitochondrial respiration was observed upon ERAP1 inhibition. Although non-mitochondrial respiration has not been adequately defined yet, several researchers have linked it to processes including nitric oxide-mediated metabolic effects^70^, cell surface oxygen consumption^71^ and peroxisomal respiration^72^. The latter might be particularly relevant here since emerging evidence suggests that peroxisomes, besides generating ROS, can also diminish them, depending on the cellular context^72^. Besides peroxisomal respiration, diminishment of ROS can also occur indirectly as a result of increased glycolysis in cancer cells^67^, also known as the Warburg effect^69^, due to diversion of substrates from the electron transport chain, which we also observed in the ERAP1 KO cells.

Although the exact mechanism is not clear, the metabolic changes described above may be related to the role of ERAP1 in peptide homeostasis in the ER. Lack of ERAP1 peptide trimming may lead to an increase in peptide concentration inside the ER, an effect that could interfere with proper protein folding. In the presence of an ER folding stressor, DTT, ERAP1 inhibition significantly affected protein folding in the ER, revealing an underlying folding problem, which was not observed in the absence of DTT, likely due to the buffering capacity of the ER environment or the sensitivity of the used method. Induction of ER stress could then be communicated to the mitochondria, given the established relationship between the two compartments, and induce metabolic changes that would help the cell adapt to the new conditions^62^. Overall, these responses may underlie the proteome changes reported here and represent a homeostatic role for ERAP1. Indeed, this aspect of ERAP1 function has been hypothesized as a pathogenetic mechanism in HLA-associated autoimmunity^60,74^.

Overall, our results support the proposed dominant role of ERAP1 as an immunopeptidome editor, while revealing secondary, but potentially, important effects in metabolic homeostasis. While the importance of these effects in cancer progression is unclear, it is possible that cancer cells may have to strike a balance between promoting immune evasion and maintaining viable cellular homeostasis. Still, the breadth of most described effects is modest, which rather highlights than negates the specialized role of ERAP1 in regulating adaptive immune responses. Since ERAP1 is a target molecule for cancer immunotherapy, it may be worth exploring whether its secondary effects on the proteome and metabolism of cancer may be exploited pharmacologically to synergize with the effects on adaptive immunity to enhance anti-tumor therapies.

## Abbreviations

ERAP1: Endoplasmic Reticulum Aminopeptidase 1
ER: Endoplasmic Reticulum
MHC-I: Major Histocompatibility Complex Class I
HLA: Human Leukocyte Antigen
ERAAP: Endoplasmic Reticulum Aminopeptidase Associated with Antigen Processing
TAP: Transporter Associated with Antigen Processing
TAP2: Transporter Associated with Antigen Presentation 2
TAPBP: Tapasin
CALR: Calreticulin
PLC: Peptide Loading Complex
MAGE: Melanoma Associated Antigen
WT: Wild-type
KO: Knock-out
PBMC: Peripheral Blood Mononuclear Cells
DMEM: Dulbecco’s Modified Eagle Medium
FBS: Fetal Bovine Serum
DIA: Data-Independent Acquisition
MBR: Match Between Runs
LC-MS/MS: Liquid chromatography-Mass Spectrometry
TFA: Trifluoroacetic Acid
ACN: Acetonitrile
FA: Formic Acid
FDR: False Discovery Rate
PCA: Principal Component Analysis
DCF: Dichlorofluorescein
ROS: Reactive Oxygen Species
RFU: Relative Fluorescence Units
Tht-T: Thioflavin T
DTT: Dithiothreitol
OCR: Oxygen Consumption Rate
ECAR: Extracellular Acidification Rate
FCCP: trifluoromethoxyphenylhydrazone

## ACKNOWLEDGMENTS

We thank Dr. Lawrence J. Stern for the generous gift of the compound 3 inhibitor used in this study.

## AUTHOR CONTRIBUTIONS

**Martha Nikopaschou:** Conceptualization, Methodology, Investigation, Formal analysis, Writing – Original Draft, Writing – Review & Editing, Visualization. **Martina Samiotaki**: Methodology, Investigation, Data Curation. **Kamila Król**: Methodology, Investigation. **Paula Gragera**: Methodology, Investigation. **Ellie Stylianaki**: Methodology, Investigation, Formal analysis. **Aroosha Raja:** Formal analysis. **Vassilis Aidinis:** Resources, Supervision. **Angeliki Chroni:** Conceptualization. **Doriana Fruci:** Resources, Supervision. **George Panayotou:** Resources, Methodology. **Efstratios Stratikos:** Conceptualization, Resources, Writing – Original Draft, Writing – Review & Editing, Visualization, Supervision, Project administration, Funding acquisition.

## DATA AVAILABILITY

All data described are available in the article and associated supporting information. This article contains supplemental data.^47,53–55,75^ Numerical values used for the generation of graphs are available upon request to the corresponding author (Efstratios Stratikos). The MS proteomics raw data have been deposited to the ProteomeXchange Consortium via the PRIDE^76^ partner repository with the dataset identifiers PXD054491 & PXD054494 (immunopeptidomics), PXD054487 (A375 proteomics) and PXD054485 (THP-1 proteomics) (http://www.ebi.ac.uk/pride/archive/).

