## Supplemental Information for "ERAP1 activity modulates the immunopeptidome but also affects the proteome, metabolism and stress responses in cancer cells"

### SUPPLEMENTAL FIGURES

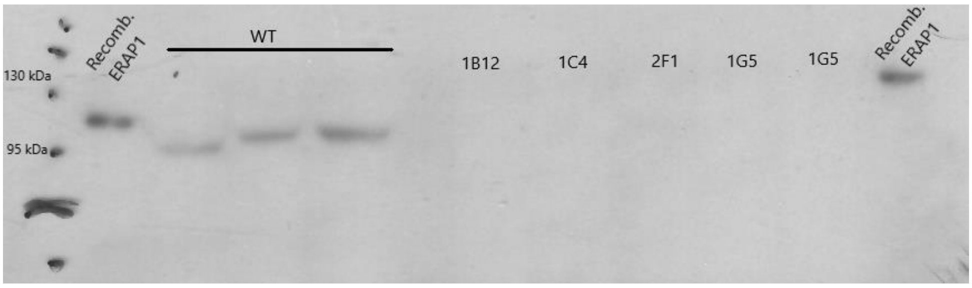


**Figure S1.** Western blot analysis showing the presence of ERAP1 in the wild-type cells and its absence in some of the obtained clones.

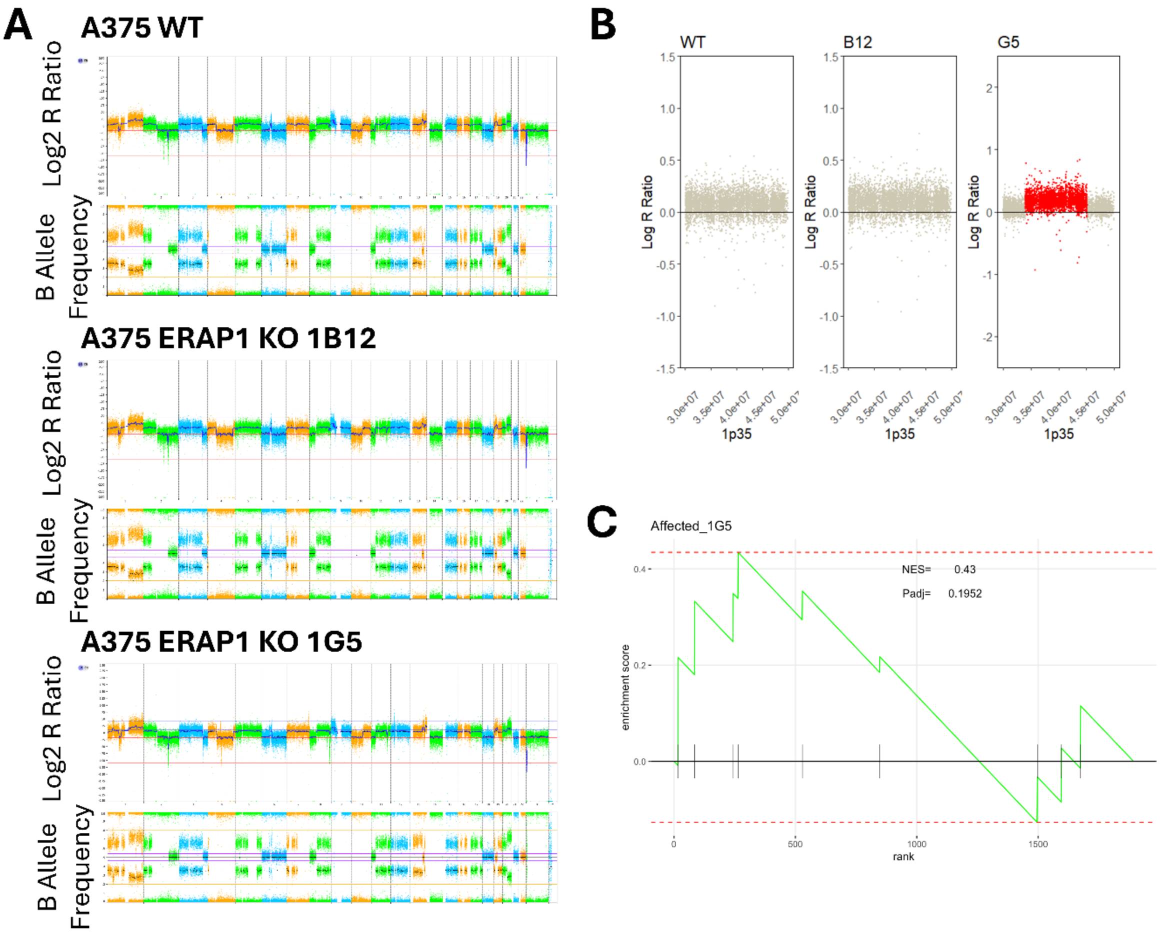


**Figure S2. A.** Whole genome copy number profiling and analysis of regions of homozygosity using SNP-arrays of the A375 cells used in this study. SNP-array based copy number profiling and analysis of regions of homozygosity using the Infinium Human CytoSNP-850K v1.2 BeadChip (Illumina, San Diego, CA, USA) showed multiple chromosomal abnormalities, as expected for cancer cells, consistent between the 3 tested clones. The only exception was the area of 1p35 for clone 1G5, in which ~20% of cells showed a partial duplication. **B.** Zoom in 1p35 area, partially affected in 1G5 cells. C. Enrichment analysis was performed with the fgsea package in R.Studio v. 4.3.3 to check whether the differentially expressed proteins in the proteomics experiment were enriched with proteins from the affected area in the 1G5 clone. The results of the analysis indicated that there is no enrichment for proteins in the affected area (Padj = 0.1952).

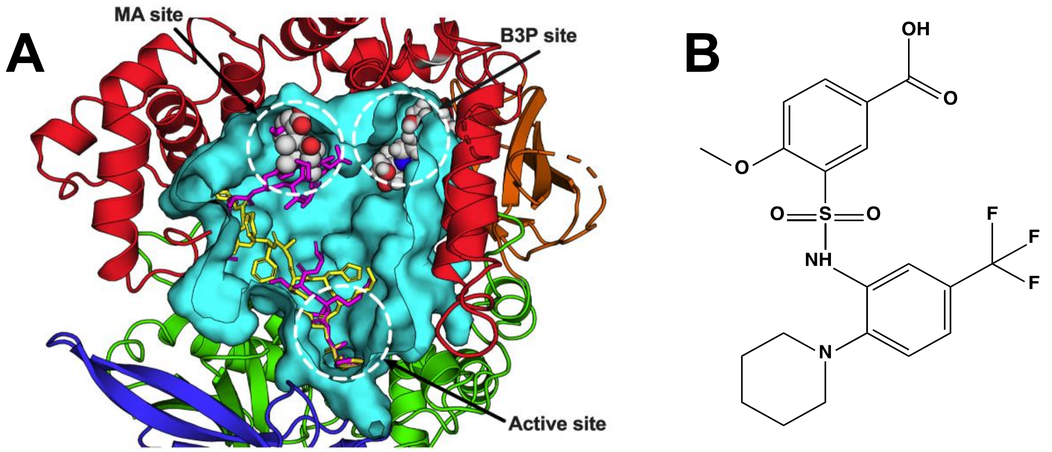


**Figure S3. A.** Schematic representation of the internal cavity of ERAP1 (in cyan cutaway view) indicating the active site of the enzyme as well as two allosteric sites found to accommodate small MW compounds in a high-resolution crystal structure (MA = malate, B3P = bis-tris-propane). Two peptide substrates crystallized with ERAP1 are shown in stick representation (10 mer peptide in yellow and 15 mer in magenta). **B.** The chemical structure of compound 3 used in this study.

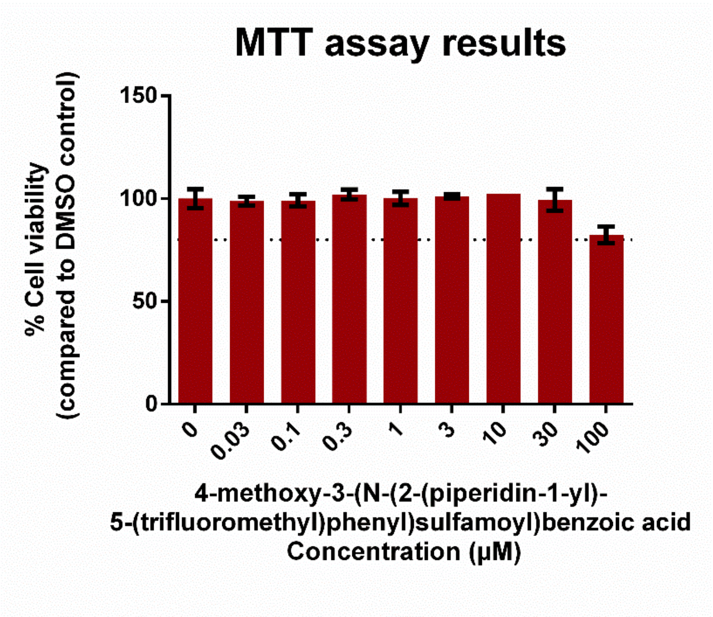


**Figure S4.** The viability of A375 cells was evaluated by the MTT assay after 48hrs exposure to different concentrations of the ERAP1 inhibitor. The dotted line at 80% indicates the threshold for adequate viability.

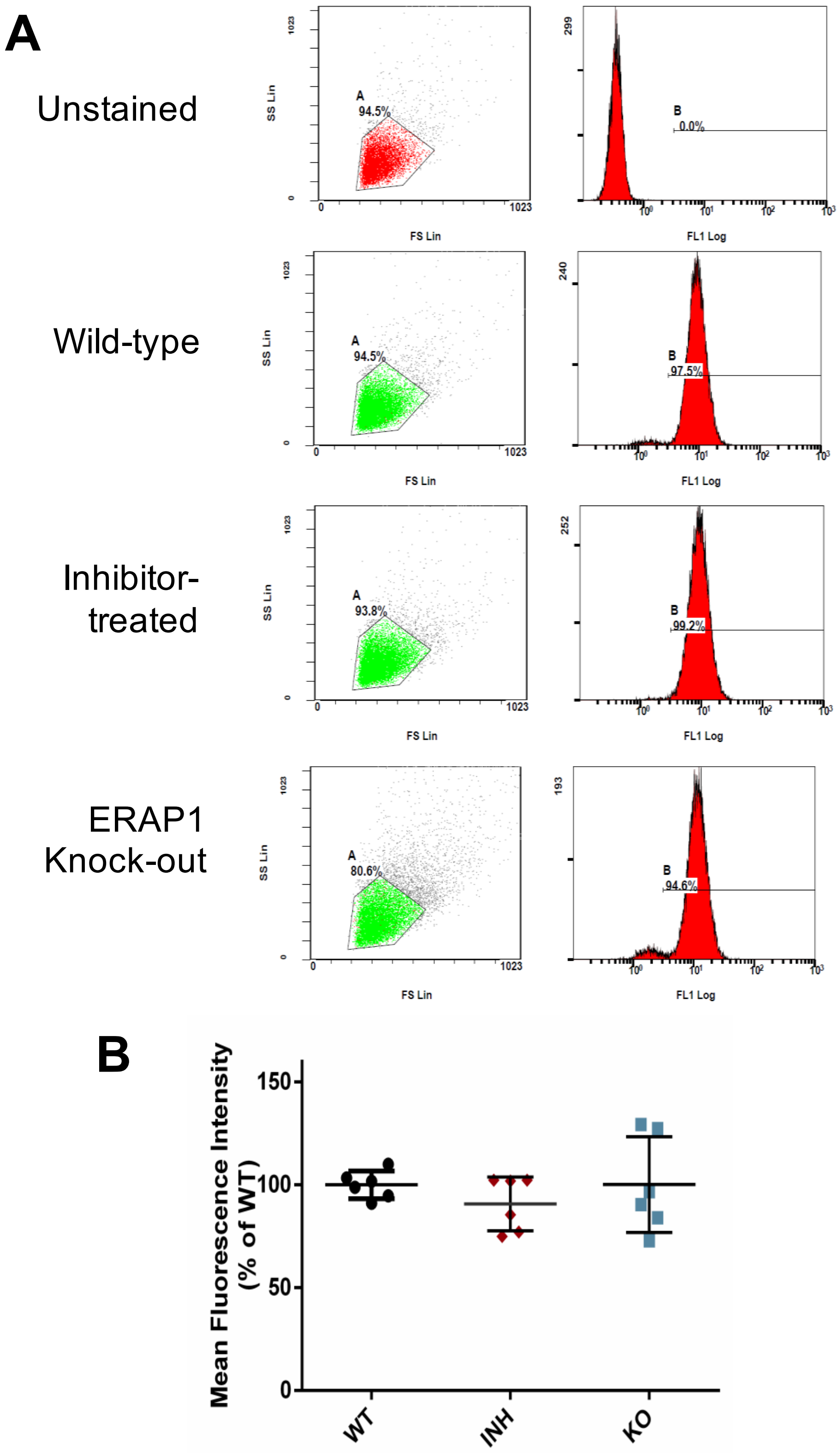


**Figure S5**: Cell-surface levels of MHC-I in A375 cells detected by FACS. **A.** Representative data from each condition indicating gating strategy and signal distribution. **B.** Normalized mean fluorescence intensity (% of Wild-type, DMSO treated cells) from 2 independent experiments with n=3 technical replicates each.

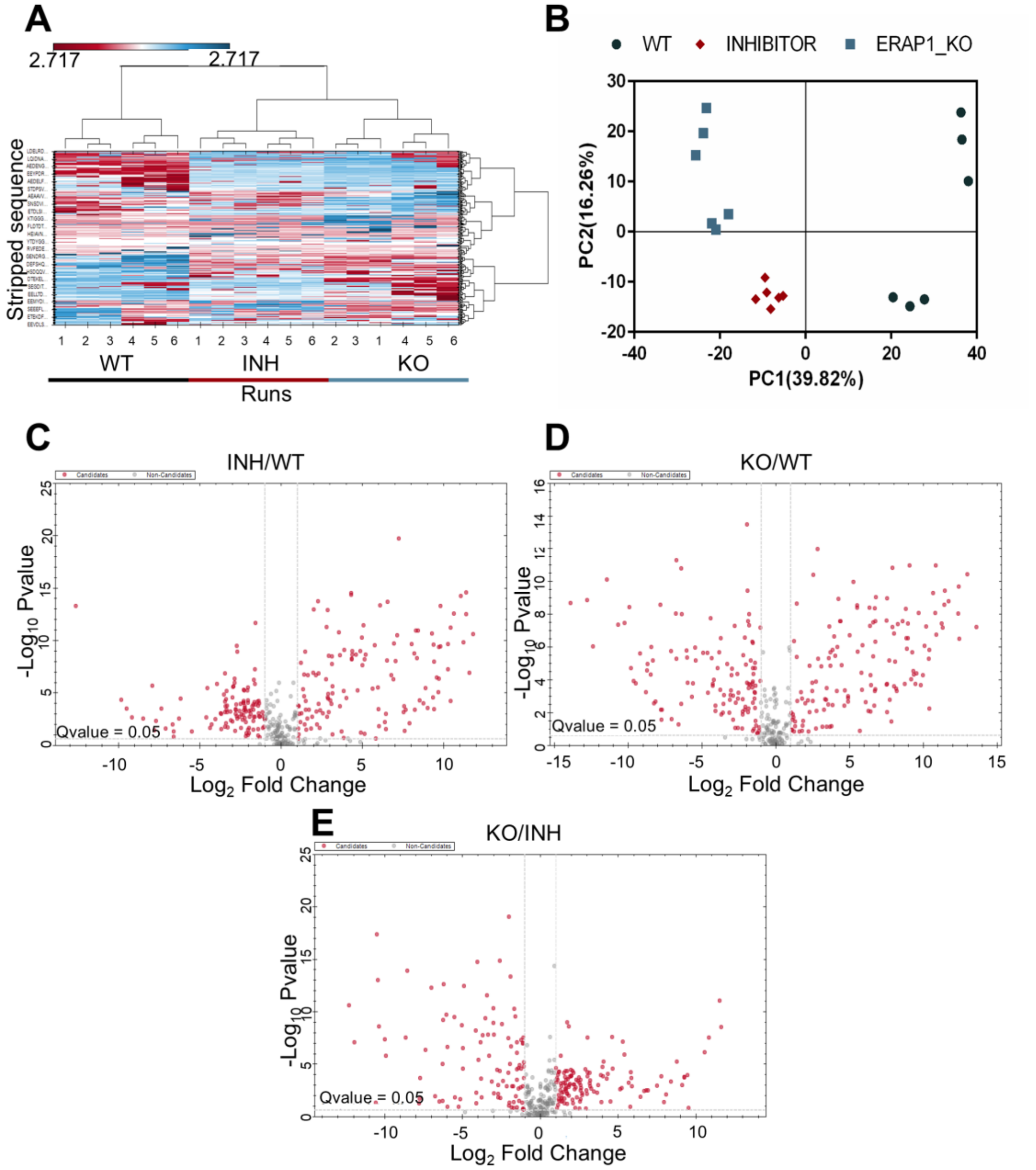


**Figure S6:** Analysis of the immunopeptidome of wild-type, inhibitor-treated & ERAP1 KO A375 cells (run 2). **A.** Heatmap from LC-MS/MS run 2 showcasing peptide distribution across different experimental conditions (bottom). Hierarchical clustering was calculated by Manhattan Distance. Colors indicate peptide intensities, ranging from low (red) to high (blue). (Graph generated with Spectronaut® v. 19) **B.** Principal Component analysis (PCA) from LC-MS/MS run 2. PCs 1 & 2 contribute to the explanation of 56.08% of sample variability. Based on these two PCs, wild-type samples (circle) inhibitor- treated samples (diamond) and KO samples (square) formed 3 distinct groups, indicating that the replicates within each group share similarities with each other but there are significant differences in the immunopeptidomes between different treatments. **C-E.** Volcano plots from LC-MS/MS run 2 indicating the statistical significance of the differences between (C) the inhibitor-treated and the wild type A375 cells, (D) the genetically modified (KO) and the wild type A375 cells and (E) the KO versus inhibitor-treated cells. Each circle represents a unique peptide sequence. Peptides with a q-value ≤0.05 and a log2 fold change ≥1 are considered statistically significant.

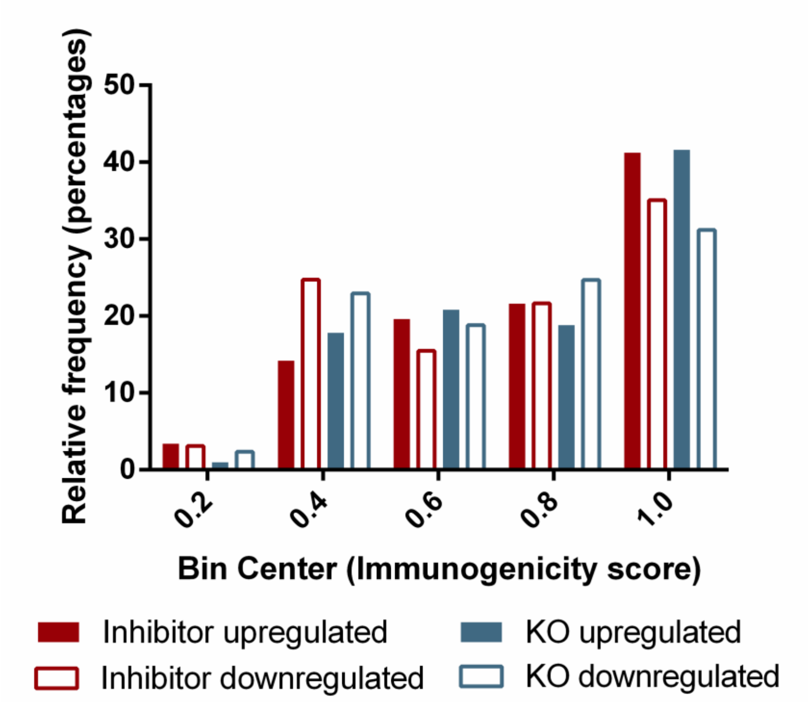


**Figure S7:** Distribution of the immunogenicity scores of the differentially expressed 9mers, obtained with the DeepImmuno algorithm.

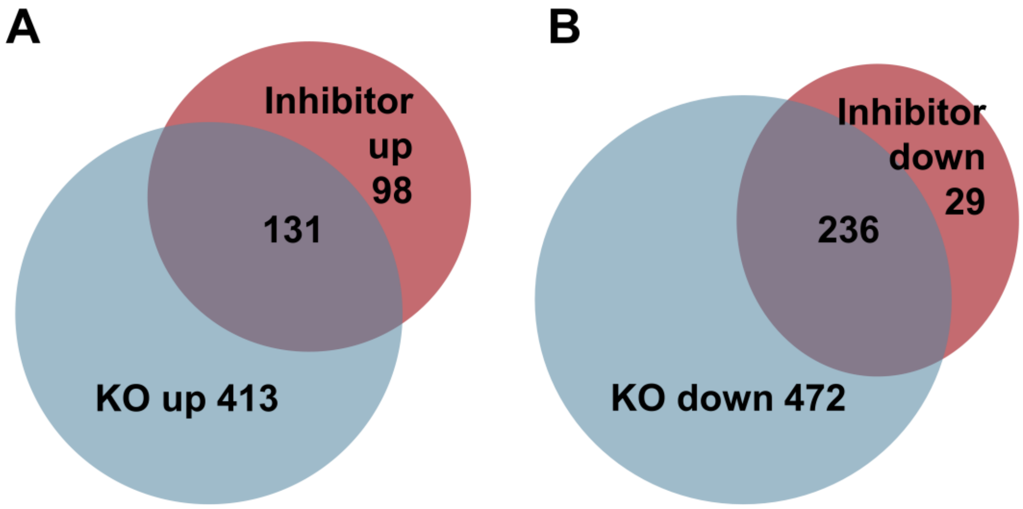


**Figure S8:** Venn diagrams, indicating the overlap of the significantly up-regulated (A) and down-regulated (B) proteins between the inhibitor-treated and the ERAP1 KO cells.

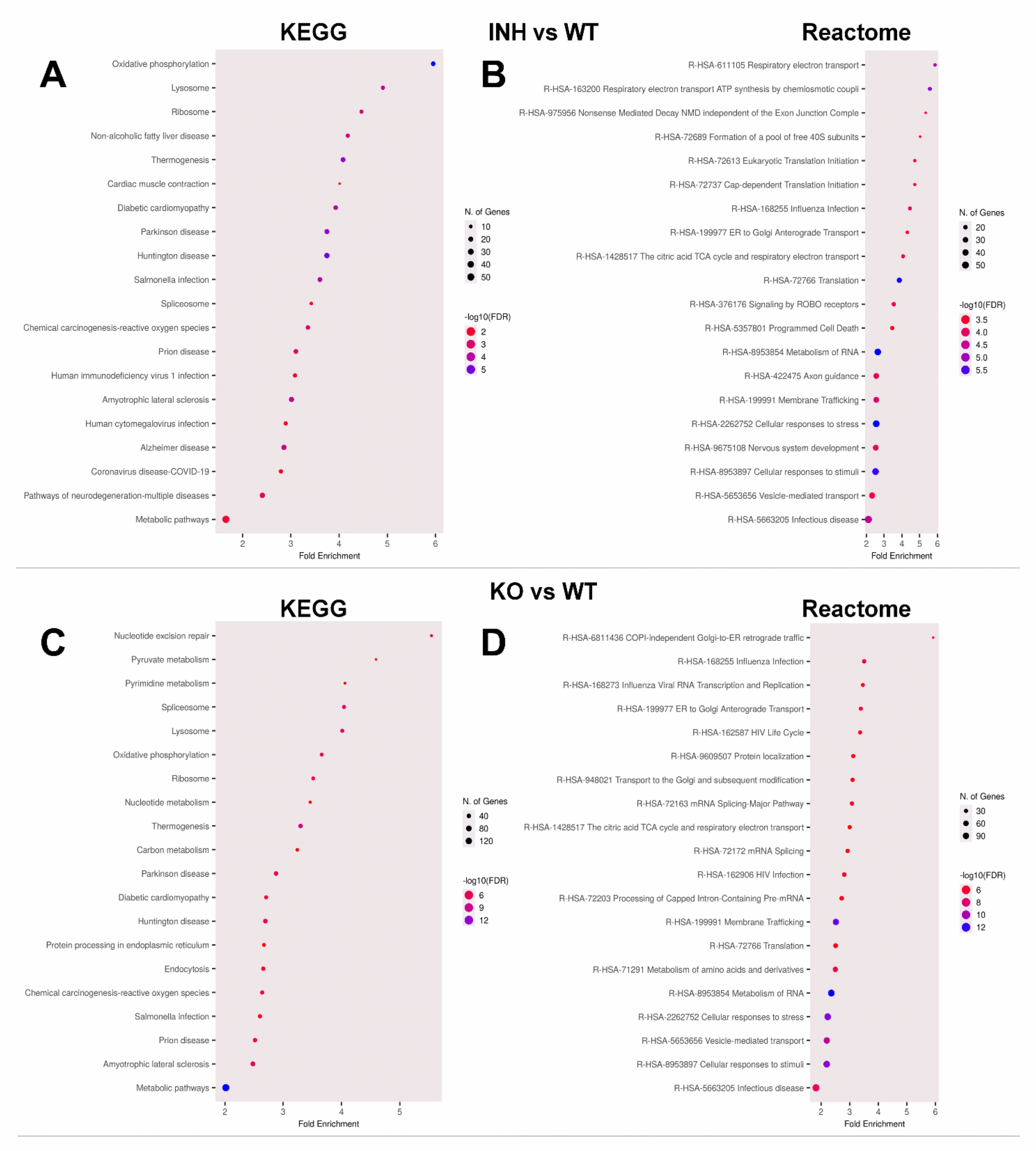


**Figure S9:** ShinyGo Pathway enrichment analysis using the KEGG and Reactome databases for A375 cells. Enriched pathways from inhibitor affected (A, B) and KO affected (C, D) proteins ordered by fold-enrichment. Colour indicates significance (FDR), while dot size indicates the number of affected genes in the given pathway.

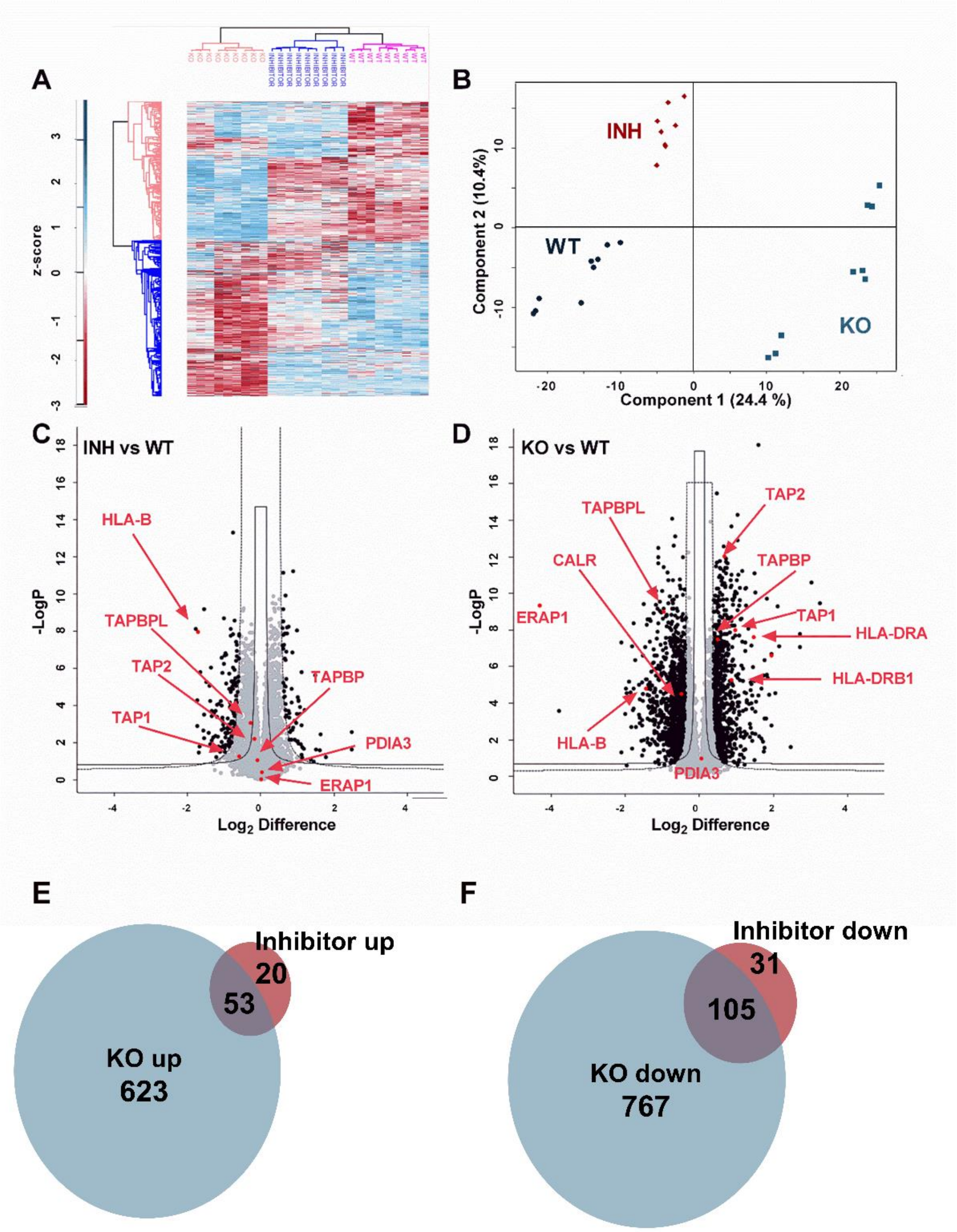


**Figure S10:** Proteomic analysis of wild-type, inhibitor-treated & ERAP1 KO THP-1 cells. **A.** Heatmap showing the distribution of proteins in the three conditions (WT, inhibitor-treated & ERAP1 KO THP-1 cells) for each replicate (three biological replicates, each measured in three technical replicates). **B.** Principal Component Analysis (PCA) of the three experimental conditions. Each point represents an injection in the LC-MS/MS. All experimental conditions can be discerned from each other. **C & D.** Hawaii plots, indicating the statistical significance of the observed differences in protein abundance between the two treatment conditions (inhibitor-treated & ERAP1 KO) and the WT cells. 209 proteins were differentially expressed in the inhibitor-treated (C) cells and 1548 proteins were differentially expressed in the KO cells (D). Select proteins that participate in antigen presentation are indicated in red. **E & F.** Venn diagrams, indicating the overlap of the significantly up-regulated (E) and down-regulated (F) proteins between the inhibitor -treated and the ERAP1 KO cells.

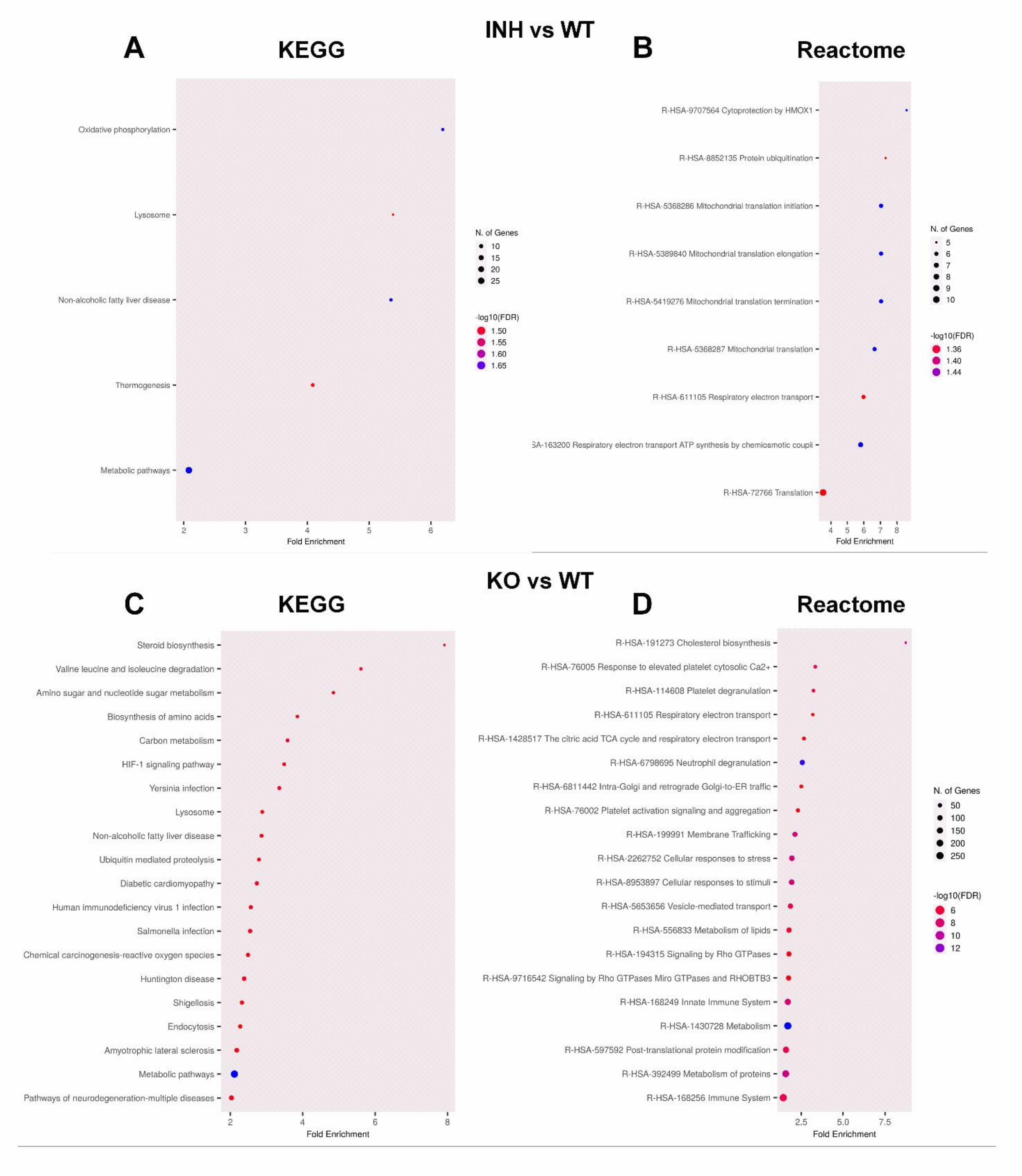


**Figure S11:** ShinyGo Pathway enrichment analysis using the KEGG and Reactome databases for THP-1 cells. Enriched pathways from inhibitor affected (A, B) and KO affected (C, D) proteins ordered by fold-enrichment. Colour indicates significance (FDR), while dot size indicates the number of affected genes in the given pathway.

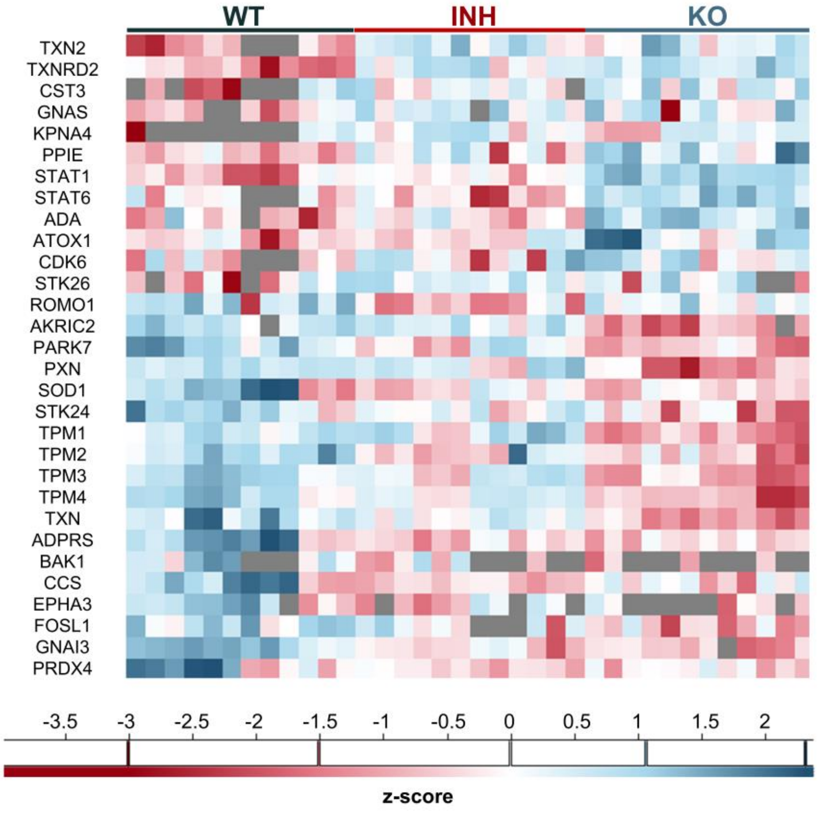


**Figure S12:** Heatmap of proteins related to response to reactive oxygen species.

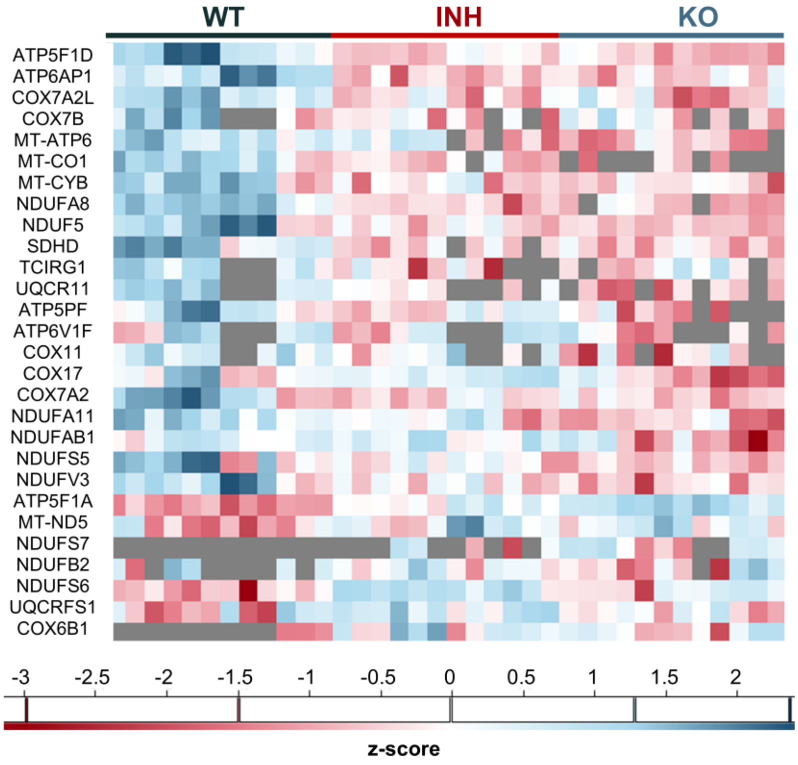


**Figure S13:** Heatmap of proteins related to oxidative phosphorylation.

**Supplemental Table S1:** Identified peptides corresponding to known MAGE antigens.

| MAGE antigens | | | |
|---|---|---|---|
| MAGE | Sequence | KO | INH |
| A3 | EVDPIGHLY | | |
| A4 | AETSYVKVL | [blue] | [blue] |
| A4 | EVDPASNTY | | |
| A4 | GSNPARYEF | [blue] | [blue] |
| A4 | KEVDPASNTY | [red] | |
| A4 | TVYGEPRKL | [blue] | [blue] |
| A11 | EVDPTSHSY | [red] | [red] |
| C2 | EEVPSGVIPNL | [blue] | [blue] |
| C2 | FVYGEPREL | [blue] | [blue] |
| [red] | Downregulated | | |
| [blue] | Upregulated | | |

**Supplemental Table S2:** Identified peptides corresponding to known antigenic epitopes In the IEDB database.

| Epitopes from IEDB | | | |
|---|---|---|---|
| Sequence | Protein Name | KO | INH |
| LLDVPTAAV | Gamma-interferon-inducible lysosomal thiol reductase precursor | [blue] | |
| KLDVGNAEV | B-cell receptor-associated protein 31 | [red] | [blue] |
| RLFDEPQLA | BTB/POZ domain-containing protein 2 | [red] | |
| TLWVDPYEV | Protein BTG1 | | |
| TMLARLASA | Chondroitin sulfate proteoglycan 4 | [red] | [blue] |
| GLIEKNIEL | DNA (cytosine-5)-methyltransferase 1 | [red] | [blue] |
| LLDLDEELRY | Chromosome 14 open reading frame 179 | [blue] | [blue] |
| EVDPIGHLY | MAGEA3 | | |
| FVYGEPREL | MAGEC2 | [blue] | [blue] |
| SIIGRLLEV | serine/threonine-protein phosphatase PP1-alpha catalytic subunit isoform 1 | | |
| IMLEALERV | Small nuclear ribonucleoprotein G (snRNP-G) (Sm protein G) | [red] | [red] |
| SEEEFLRTY | Beta-catenin-interacting protein 1 | [red] | |
| [red] | Downregulated | | |
| [blue] | Upregulated | | |

**Supplemental Table S3:** Identified peptides originating from unannotated proteins.

| Unannotated proteins | | | | |
|---|---|---|---|---|
| Sequence | Length | PG.ProteinGroups | KO | INH |
| AEEPLAGRTW | 10 | GN=POLD4 3' Overlap dORF | | |
| ALAAVVTEV | 9 | GN=DDX3X 3' dORF | | |
| EEAVVLRGL | 9 | GN=WBP1 5' uORF | | |
| EEYRGGNNW | 9 | GN=TTC37 5' uORF | | |
| FSNDALKTY | 9 | GN=FAM127B 3' Overlap dORF | | |
| IEVDGGRDW | 9 | GN=SLC39A7 5' uORF | | |
| IFSQRSYSY | 9 | GN=NFAT5 Out-of-Frame | | |
| ISDPGVQGY | 9 | GN=IKBKAP 5' Overlap uORF | | |
| LVSNGVLVV | 9 | GN=PLA2G4B 3' Overlap dORF | | |
| RTAAAAAGAASW | 12 | GN=URI1 5' uORF | | |
| RTDQFYVVY | 9 | GN=PTGES3 ncRNA Processed Transcript | | |
| SAAWDRPPL | 9 | GN=MPG Out-of-Frame | | |
| STSTILRSF | 9 | GN=GLCE ncRNA Processed Transcript | | |
| TDGVSLLLP | 9 | GN=TFB1M 5' Overlap uORF | | |
| TDNRTDIFY | 9 | GN=nan lincRNA | | |
| VEDPIAEGGR | 10 | GN=RUBCN Truncated | | |
| VTEKVYADTGLY | 12 | GN=TMEM168 5' uORF | | |

Downregulated

Upregulated

### SUPPLEMENTAL METHODS

#### Isolation of genomic DNA

A375 wild type and two clones of ERAP1 silenced cells (1G5 and 1B12) were washed twice with ice-cold PBS and incubated overnight at 37 °C with lysis buffer (10 mM NaCl, 10 mM Tris–HCl pH 7.5, 10 mM EDTA, 0.5% SDS and 100 µg/ml RNAse A and 100 µg/ml protein- ase K). Genomic DNA was isolated by phenol/$CHCl_3$ extraction and precipitated with 1/10 vol of 3 M sodium acetate, pH 5.2, and 2 vol of 100% ethanol. The pellet was washed with 70% ethanol, re-suspended in sterile milli-Q $H_2O$ and quantitated based on absorption at 260/280 nm.

#### High-density SNP-array analysis

Genomic DNA from A375 wild type and ERAP1-silenced cells was used for SNP-array copy number profiling and analysis of regions of homozygosity with the Infinium Human CytoSNP-850K v1.2 BeadChip (Illumina, San Diego, CA, USA). This array has approximately 850,000 single nucleotide polymorphism (SNP) markers spanning the genome and can detect genomic insertions, deletions, and regions of homozygosity. Data analysis was performed with NxClinical software v6.0 (Bionano genomics, San Diego, CA, USA). Human genome build Feb. 2009 GRCh37/hg19 was used and results were classified with BENCH Lab CNV software (Agilent, Santa Clara, CA, USA).

#### MTT cytotoxicity assay

Wild-type A375 cells were seeded in a 96-well plate at a density of 50,000 cells/ml. 24 hours later, cells were treated with the inhibitor at a concentration range between 0 and 100 µM in the presence of 0.5% DMSO. Cells were incubated for 48 hours at 37 °C, 5% $CO_2$. The medium was replaced with 100 µl fresh medium containing 1 mg/ml 3-(4,5-Dimethylthiazol-2-yl)-2,5-Diphenyltetrazolium Bromide (MTT) and the plate was incubated for 4 hours. MTT containing medium was removed, 100 µl dimethylsulfoxide (DMSO) per well were added and the plate was shaken for 15 minutes prior to measurement of absorbance at 540 and 620 nm.

#### Surface MHC I expression

On treatment day 6, ca. $10^6$ cells were detached from the wells of a 12-well plate with 10 mM Ethylenediamine tetraacetic acid (EDTA) in Phosphate-buffered saline (PBS). Cells were washed with PBS and FACS buffer (1% Bovine serum albumin, 0.02% Sodium azide in PBS) prior to incubation with the Mouse, anti-human HLA-ABC FITC labelled antibody, diluted 1:25 (Biorad, MCA81F). A negative control was also prepared by incubating with FACS buffer without the addition of the antibody. The incubation took place for 30 minutes on ice and the cells were protected from light. Subsequently, cells were washed twice with FACS buffer, resuspended in 500 µl FACS buffer- 1% Formic Acid 1:1 and transferred to FACS tubes for the analysis with a Cytomics™ FC 500 (Beckman Coulter) cytometer working with the CXP Analysis software (v 2.2.).

#### Spectronaut settings

Settings Used: BGS Factory Settings

├─ DIA Analysis\Calibration

| ├─ MZ Extraction Strategy: Maximum Intensity

| ├─ Precision iRT: True

| | ├─ Exclude De-amidated Peptides: True

| | └─ iRT <-> RT Regression Type:Local (Non-Linear) Regression

| ├─ MS1 Mass Tolerance Strategy: System Default
| └─ MS2 Mass Tolerance Strategy:System Default
├─ DIA Analysis\Identification
| ├─ Precursor Qvalue Cutoff: 0.01
| ├─ Precursor PEP Cutoff: 0.2
| ├─ Protein Qvalue Cutoff (Experiment): 0.01
| ├─ Protein Qvalue Cutoff (Run): 0.05
| ├─ Protein PEP Cutoff: 0.75
| ├─ Single Hit Definition: By Stripped Sequence
| ├─ Exclude Single Hit Proteins: False
| ├─ Exclude Duplicate Assays: True
| ├─ Exclude Predicted Fragment Scores: False
| ├─ Generate Decoys: True
| | ├─ Decoy Generation Method: Mutated
| | | └─ Preferred Fragment Source: NN Predicted Fragments
| | └─ Decoy Limit Strategy: Dynamic
| | └─ Library Size Fraction: 0.1
| └─ Pvalue Estimator: Kernel Density Estimator
├─ DIA Analysis\Pipeline Mode
| ├─ Export All XICs: False
| ├─ Generate SNE File: True
| | └─ Store Ion traces in SNE: True
| ├─ Post Analysis Reports:
| | ├─ Binned CVs:False
| | ├─ Binned Identification: False
| | ├─ CV Density Line Chart: False
| | ├─ CVs Below X Bar Chart: False
| | ├─ Data Completeness Bar Chart: False
| | ├─ Run Identifications Bar Chart: False
| | └─ Scoring Histograms:False
| ├─ Report Schema: BGS Factory Report (Normal)
| └─ Reporting Unit: Across Experiment
├─ DIA Analysis\Post Analysis
| ├─ Differential Abundance Testing: Unpaired t-test

```
|  |  ├─ Assume Equal Variance:    False
|  |  ├─ Group-Wise Testing Correction:    False
|  |  ├─ Log2 Ratio Candidate Filter:    0.58
|  |  └─ Confidence Candidate Filter:    Qvalue
|  |     └─ Confidence:    0.05
|  ├─ Differential Abundance Grouping:    Minor Group (Quantification Settings)
|  |  └─ Smallest Quantitative Unit:Minor Group (Quantification Settings)
|  |     └─ Use All MS-Level Quantities:    True
|  ├─ Calculate Explained TIC:    None
|  ├─ Calculate Sample Correlation Matrix:False
|  └─ Hierarchical Clustering:    True
|     ├─ Distance Metric:    Manhattan Distance
|     ├─ Linkage Strategy:    Ward's Method
|     ├─ Order Runs by Clustering:    True
|     └─ Z-score Transformation:    True
├─ DIA Analysis\Protein Inference
|  └─ Protein Inference Workflow:    Automatic
|     └─ Inference Algorithm:IDPicker
├─ DIA Analysis\PTM Workflow
|  ├─ [Beta] Input Normalization Strategy: None
|  └─ PTM Localization:    False
├─ DIA Analysis\Quantification
|  ├─ Precursor Filtering:    Identified (Qvalue)
|  |  ├─ Imputation Strategy:    None
|  |  └─ Multi Channel Qvalue Filter:    Group Qvalue
|  ├─ Proteotypicity Filter:    None
|  ├─ Protein LFQ Method:    Automatic
|  ├─ Quantity MS Level:    MS2
|  ├─ Quantity Type:    Area
|  ├─ Cross-Run Normalization:    True
|  |  ├─ Normalization Filter Type: None
|  |  ├─ Normalization Strategy:    Automatic
|  |  └─ Row Selection:    Automatic
|  |     └─ Multi Channel Qvalue Filter:    Group Qvalue
```

| ├─ Quantification window: Synchronized
| ├─ Interference Correction: True
| | ├─ Only Identified Peptides: True
| | ├─ Exclude All Multi-Channel Interferences: True
| | ├─ MS1 Min: 2
| | └─ MS2 Min: 3
| ├─ Major (Protein) Grouping: by Protein Group Id
| ├─ Minor (Peptide) Grouping: by Stripped Sequence
| ├─ Major Group Quantity: Mean peptide quantity
| ├─ Major Group Top N: True
| | ├─ Max: 3
| | └─ Min: 1
| ├─ Minor Group Quantity: Mean precursor quantity
| ├─ Minor Group Top N: True
| | ├─ Max: 3
| | └─ Min: 1
| └─ DeepQuant Correction [Beta]: False
├─ DIA Analysis\Workflow
| ├─ Method Evaluation: False
| ├─ Profiling Strategy: None
| ├─ Run Limit for directDIA Library: -1
| ├─ Hybrid (DDA + DIA) Library: False
| └─ Unify Peptide Peaks Strategy: None
├─ DIA Analysis\XIC Extraction
| ├─ XIC IM Extraction Window: Dynamic
| | └─ Correction Factor: 1
| ├─ XIC RT Extraction Window: Dynamic
| | └─ Correction Factor: 1
| ├─ MS1 Mass Tolerance Strategy: Dynamic
| | └─ Correction Factor: 1
| └─ MS2 Mass Tolerance Strategy: Dynamic
| └─ Correction Factor: 1
├─ Pulsar Search\Identification
| ├─ PSM FDR: 0.05

```
|  ├─ Peptide FDR:  0.05
|  ├─ Protein Group FDR:    1
|  ├─ directDIA Workflow:  directDIA+ (Deep)
|  |  └─ RT Sampling Reduction:     1
|  └─ PTM Localization Filter:        False
├─ Pulsar Search\Labeling
|  └─ Channels:
|     ├─ Channel 1:  False
|     ├─ Channel 2:  False
|     └─ Channel 3:  False
├─ Pulsar Search\Modifications
|  ├─ Max Variable Modifications: 5
|  └─ Select Modifications:
|     ├─ Fixed Modifications::
|     └─ Variable Modifications: :    Acetyl (Protein N-term), Oxidation (M)
├─ Pulsar Search\Peptides
|  ├─ Enzymes / Cleavage Rules:
|  ├─ Digest Type:  Unspecific
|  ├─ Max Peptide Length: 25
|  ├─ Min Peptide Length:  7
|  ├─ Missed Cleavages:     2
|  └─ Toggle N-terminal M: True
├─ Pulsar Search\Result Filters
|  ├─ Fragment Ions:
|  |  ├─ Ion AA Length:       True
|  |  |  └─ N:3
|  |  ├─ Ion Charge: False
|  |  ├─ Ion Loss Type:       False
|  |  ├─ Ion Type:    False
|  |  ├─ m/z :        True
|  |  |  ├─ Max:      3000
|  |  |  └─ Min:      200
|  |  ├─ Overlapping between Channels:   False
|  |  └─ Relative Intensity:  True
```

```
| |    └─ Min:        1
| └─ Precursors:
|    ├─ Amino Acids:          False
|    ├─ Best N Fragments per Peptide:      True
|    | ├─ Max:       6
|    | └─ Min:       3
|    ├─ Best N Peptides per Protein Group:False
|    ├─ Channel Count:       False
|    ├─ FASTA Matched:       False
|    ├─ Missed Cleavage:   False
|    ├─ Modifications:       None
|    ├─ Peptide Charge:      False
|    └─ Proteotypicity:       False
├─ Pulsar Search\Speed-Up
| ├─ MS2 Index:    Automatic
| └─ diaPASEF Pre-Processing:       Fast (Spectronaut 19)
├─ Pulsar Search\Tolerances
| └─ Tolerance Parameters:
|    ├─ Thermo IonTrap:
|    | ├─ Calibration Search:         Dynamic
|    | | ├─ MS1 Correction Factor:1
|    | | └─ MS2 Correction Factor:1
|    | └─ Main Search:        Dynamic
|    |    ├─ MS1 Correction Factor:1
|    |    └─ MS2 Correction Factor: 1
|    ├─ Thermo Orbitrap:
|    | ├─ Calibration Search:         Dynamic
|    | | ├─ MS1 Correction Factor:1
|    | | └─ MS2 Correction Factor:1
|    | └─ Main Search:        Dynamic
|    |    ├─ MS1 Correction Factor:1
|    |    └─ MS2 Correction Factor: 1
|    └─ TOF:
|       ├─ Calibration Search:         Dynamic
```

```
|       |  ├─ MS1 Correction Factor:1
|       |  └─ MS2 Correction Factor: 1
|       └─ Main Search:         Dynamic
|          ├─ MS1 Correction Factor: 1
|          └─ MS2 Correction Factor:  1
└─ Pulsar Search\Workflow
   ├─ Fragment Ion Selection Strategy:       Intensity Based
   ├─ In-Silico Generate Missing Channels: False
   └─ Use DNN Predicted Ion Mobility:        Auto
[END-SETTINGS]
```

**DIANN settings**

DIA-NN 1.8 (Data-Independent Acquisition by Neural Networks)
Compiled on Jun 28 2021 14:55:31
Current date and time: Mon Jul 4 09:47:37 2022
CPU: GenuineIntel Intel(R) Core(TM) i9-10900 CPU @ 2.80GHz
SIMD instructions: AVX AVX2 FMA SSE4.1 SSE4.2
Logical CPU cores: 20
diann.exe --f Y:\HF_X_2022…
--lib --threads 20 --verbose 1 --out Y:\….tsv --qvalue 0.01 --matrices --out-lib Y:\….tsv --gen-spec-lib --predictor --fasta D:\Fasta\uniprot_human_reviewed_110422.fasta --fasta-search --min-fr-mz 200 --max-fr-mz 1800 --met-excision --cut K*,R* --missed-cleavages 2 --min-pep-len 7 --max-pep-len 30 --min-pr-mz 300 --max-pr-mz 1500 --min-pr-charge 2 --max-pr-charge 4 --var-mods 3 --var-mod UniMod:35,15.994915,M --double-search --individual-mass-acc --individual-windows --reanalyse --smart-profiling --peak-center

Thread number set to 20
Output will be filtered at 0.01 FDR
Precursor/protein x samples expression level matrices will be saved along with the main report
A spectral library will be generated
Deep learning will be used to generate a new in silico spectral library from peptides provided
Library-free search enabled
Min fragment m/z set to 200
Max fragment m/z set to 1800
N-terminal methionine excision enabled
In silico digest will involve cuts at K*,R*
Maximum number of missed cleavages set to 2
Min peptide length set to 7
Max peptide length set to 30
Min precursor m/z set to 300
Max precursor m/z set to 1500
Min precursor charge set to 2
Max precursor charge set to 4

Maximum number of variable modifications set to 3
Modification UniMod:35 with mass delta 15.9949 at M will be considered as variable
Neural networks will be used for peak selection
Mass accuracy will be determined separately for different runs
Scan windows will be inferred separately for different runs
A spectral library will be created from the DIA runs and used to reanalyse them; .quant files will only be saved to disk during the first step
When generating a spectral library, in silico predicted spectra will be retained if deemed more reliable than experimental ones
Fixed-width center of each elution peak will be used for quantification
DIA-NN will optimise the mass accuracy separately for each run in the experiment. This is useful primarily for quick initial analyses, when it is not yet known which mass accuracy setting works best for a particular acquisition scheme.
